# Neuropsychiatric Phenotypes and a Distinct Constellation of ASD Features in 3q29 Deletion Syndrome: Results from the 3q29 Registry

**DOI:** 10.1101/386243

**Authors:** Rebecca M Pollak, Melissa M Murphy, Michael P Epstein, Michael E Zwick, Cheryl Klaiman, Celine A Saulnier, the Emory 3q29 Project, Jennifer G Mulle

## Abstract

**Background:** The 1.6 Mb 3q29 deletion is associated with neurodevelopmental and psychiatric phenotypes, including increased risk for autism spectrum disorder (ASD) and a 20-40-fold increased risk for schizophrenia. However, the phenotypic spectrum of the deletion, particularly with respect to ASD, remains poorly described.

**Methods:** We ascertained individuals with 3q29 deletion syndrome (3q29Del, “cases”, n=93, 58.1% male) and typically developing controls (n=64, 51.6% male) through the 3q29 registry (https://3q29deletion.patientcrossroads.org). Self-report of neuropsychiatric illness was evaluated for 93 cases. Subsets of participants were evaluated with the Social Responsiveness Scale (SRS, n=48 cases, 56 controls), Social Communication Questionnaire (SCQ, n=33 cases, 46 controls), Autism Spectrum Screening Questionnaire (ASSQ, n=24 cases, 35 controls), and Achenbach Behavior Checklists (n=48 cases, 57 controls).

**Results:** 3q29Del cases report a higher prevalence of autism diagnoses versus the general population (29.0% vs. 1.47%, p<2.2E-16). Notably, 3q29 deletion confers a greater influence on risk for ASD in females (OR=41.8, p=4.78E-05) than in males (OR=24.6, p=6.06E-09); this is aligned with the reduced male:female bias from 4:1 in the general population to 2:1 in our study sample. Although 71% of cases do not report a diagnosis of ASD, there is evidence of significant social disability (3q29Del SRS *T-score*=71.8, control SRS *T-score*=45.9, p=2.16E-13). Cases also report increased frequency of generalized anxiety disorder compared to controls (28.0% vs. 6.2%, p=0.001), which is mirrored by elevated mean scores on the Achenbach DSM-oriented sub-scales (p<0.001). Finally, cases show a distinct constellation of ASD features on the SRS as compared to idiopathic ASD, with substantially elevated Restricted Interests and Repetitive Behaviors, but only mild impairment in Social Motivation.

**Conclusions:** Our sample of 3q29Del is significantly enriched for ASD diagnosis, especially among females, and features of autism may be present even when an ASD diagnosis is not reported. Further, the constellation of ASD features in this population is distinct from idiopathic ASD, with substantially less impaired social motivation. Our study implies that ASD evaluation should be the standard of care for individuals with 3q29Del. From a research perspective, the distinct ASD subtype present in 3q29Del is an ideal entry point for expanding understanding of ASD.

## BACKGROUND

3q29 deletion syndrome (3q29Del) is a rare (∼1 in 30,000) [1, 2] genomic disorder characterized by a 1.6 Mb typically *de novo* deletion on chromosome 3 [3–5]. The interval contains 21 distinct protein-coding genes, 3 antisense transcripts, 1 long noncoding RNA, and 1 microRNA. Our understanding of the syndrome phenotype continues to evolve. Initial reports found developmental delay/intellectual disability universal among 3q29 deletion carriers, though some case reports have since identified individuals without cognitive impairment [6]. The 3q29 deletion is associated with a 20-40-fold increased risk for schizophrenia (SZ), with multiple replication studies supporting this association [7–11]. Case reports also indicate other neuropsychiatric phenotypes may exist, including attention deficit/hyperactivity disorder (ADHD) and bipolar disorder [3, 4, 12–16]. Previous work by our team examining self-report data from 44 individuals with 3q29Del revealed a high prevalence (∼20%) of generalized anxiety disorder [5]. Further, case reports have long suggested an association with autism spectrum disorder (ASD), and studies with large sample sizes indicate that the 3q29 deletion may confer a 19-fold increased risk for ASD (p = 0.001) [17, 18].

The range of neuropsychiatric manifestations in 3q29Del is consistent with other genomic disorders. For example, the 22q11.2 deletion has a well-known association with schizophrenia but is also associated with intellectual disability (ID), ASD, anxiety, mood disorders, and ADHD [19, 20]. A similar constellation of phenotypes, including ASD, ADHD, ID, SZ, and anxiety, has been identified in 16p11.2 deletion and duplication syndromes [21, 22], 7q11.23 duplication syndrome [23], and 1q21.1 deletion syndrome [24]. Thus, risk for multiple neuropsychiatric phenotypes appears to be a feature common to many genomic disorders, including 3q29 deletion syndrome.

The present study aims to improve the current understanding of 3q29 deletion-associated neuropsychiatric and neurodevelopmental phenotypes, and ASD in particular, by examining data from comprehensive, standardized questionnaires in the largest cohort of individuals with 3q29Del ever assembled. Developing a clearer and more comprehensive picture of 3q29 deletion-associated phenotypes will aid in management of the syndrome for both families and clinicians, which may in turn improve long-term outcomes. Additionally, a careful description of the phenotypic spectrum of 3q29Del provides a basis for cross-disorder comparison between genomic disorders, which may ultimately create inroads for identifying common mechanisms underlying 3q29Del and similar CNV disorders.

## METHODS AND MATERIALS

### Sample

Individuals with 3q29Del were ascertained through the internet-based 3q29 deletion registry (https://3q29deletion.patientcrossroads.org) as previously reported [5]. Briefly, information about the registry was emailed to health care providers, medical geneticists, genetic counselors, and support organizations; the registry is also advertised via Google AdWords, where specific keywords were chosen to target the registry website in internet searches.

Participant recruitment, informed consent and assent, and data collection are all performed through the registry website. Data were securely downloaded and de-identified for analysis. After data cleaning of the electronic records (removing spam accounts, duplicate records, and related individuals), 93 3q29Del registrants (58.1% male) were included in the present study, ranging in age from 0.1-41.0 years (mean = 10.0±8.6 years). Clinical diagnosis of 3q29 deletion syndrome was confirmed in 58% of our study subjects via review of clinical genetics reports and/or medical records. To confirm that adaptation of standardized questionnaires to an online format did not skew results, 64 typically developing controls (51.6% male) were included, ranging in age from 1.0-41.0 years (mean = 9.9±7.2 years). Controls were recruited via emails sent to intramural CDC and Emory listservs and invited to fill out surveys in an identical fashion to cases. Controls reporting a clinical diagnosis of any neurodevelopmental disorder were excluded (n = 1). Description of the study sample can be found in Table 1. This study was approved by Emory University’s Institutional Review Board (IRB00064133).

**Table 1:** Characteristics of study participants with 3q29Del and controls.

|  | 3q29 Deletion Syndrome | Control | P value |
| --- | --- | --- | --- |
| Age, years (mean ± SD) | 10.0 ± 8.6 | 9.9 ± 7.2 | 0.945 |
| Sex (n, %) |  |  | 0.521 |
| Male | 54 (58.1%) | 33 (51.6%) |  |
| Female | 39 (41.9%) | 31 (48.4%) |  |
| Race (n, %) |  |  | 0.0003 |
| White | 81 (87.1%) | 41 (64.1%) |  |
| Black/African American | 2 (2.2%) | 12 (18.8%) |  |
| Other | 10 (10.8%) | 9 (14.1%) |  |
| Blank | 0 (0%) | 2 (3.1%) |  |
| Heart Defect (n, %) |  |  | 2.37E-07 |
| Yes | 27 (29.0%) | 2 (3.1%) |  |
| No | 54 (58.1%) | 61 (95.3%) |  |
| Blank | 12 (12.9%) | 1 (1.6%) |  |
| Age at Walking (n, %) |  |  | 2.16E-09 |
| Normal | 42 (45.7%) | 60 (93.8%) |  |
| Delayed | 23 (25.0%) | 1 (1.6%) |  |
| Extremely Delayed | 12 (13.0%) | 1 (1.6%) |  |
| Unsure | 10 (10.9%) | 2 (3.1%) |  |
| Not applicable | 5 (5.4%) | 0 (0%) |  |

Demographic data collected from the custom Medical & Demographic Questionnaire completed by participants upon enrollment in the online 3q29 Registry. P values were calculated with
Student's t-test (age), Fisher's exact test (race, heart defect, age at walking), or Pearson's chi square test (sex).

### Questionnaires

Upon registration, the participant or his/her parent completed a custom medical and demographic questionnaire. This questionnaire includes questions on the sex, birthdate, race, and ethnicity of the participant, as well as a detailed medical history, including developmental milestones and prior clinical diagnoses of any neuropsychiatric or neurodevelopmental disorders [5].

Four standardized questionnaires were used to assess ASD-related symptomology and general behavioral problems in the participants. The Social Responsiveness Scale (SRS; preschool, school-age, and adult forms; n = 48 3q29Del, 56 controls) is a 65-item, 4 point Likert-scaled questionnaire designed to assess ASD-related symptoms along a normative continuum [25]. The Social Communication Questionnaire (SCQ, n = 33 3q29Del, 46 controls) is a 40-item, yes/no questionnaire designed to assess ASD-related symptoms keyed to DSM criteria [26]. The Autism Spectrum Screening Questionnaire (ASSQ, n = 24 3q29Del, 35 controls) is a 27-item, yes/somewhat/no questionnaire designed to assess ASD-related symptoms in high-functioning individuals with no to mild ID [27]. The Child Behavior Checklist (CBCL) and Adult Behavior Checklist (ABCL) are 100-, 113-, or 126-item (CBCL preschool, CBCL school-age, and ABCL, respectively; n = 48 3q29Del, 57 controls), 3 point Likert-scaled questionnaires designed to assess behavioral or developmental problems [28, 29]. Data from the CBCL and ABCL were pooled for analysis. All standardized questionnaires were adapted for the online 3q29 deletion registry and were completed by the participant or parent/guardian of the participant upon registration. Some participants were not eligible to complete the standardized questionnaires because the proband was too young. Demographic characteristics of the respondents for each questionnaire can be found in Table S1, demonstrating that the average age and sex distribution of participants who completed the medical and demographic questionnaire was not different from the average age and sex distribution of participants who completed each standardized form.

### Analysis

Data from standardized questionnaires were imported into R [30] and were recoded and scored according to the publisher’s guidelines. Features of interest from the medical history questionnaire (heart defects, age at walking, ASD diagnosis, global developmental delay/mental retardation (GDD/MR) diagnosis) were recoded for analysis as follows: heart defects, yes/no; age at walking, binned as normal (≤18 months), delayed (19-24 months), and extremely delayed (>24 months); ASD diagnosis, yes/no; GDD/MR diagnosis yes (reported diagnosis of global developmental delay and/or mental retardation)/no. To compare responses between 3q29Del cases and controls, linear models and logistic regression models were implemented using the stats R package [30]. To perform case-only analysis within 3q29Del cases, linear models and logistic regression models were implemented using the stats R package [30] and cumulative link proportional-odds models were implemented using the ordinal R package [31]. All statistical models included age, race, and sex as covariates. To compare rates of self-reported diagnoses and demographic parameters between 3q29Del cases and controls, Fisher’s exact test was implemented using the stats R package [30]. To compare rates of self-reported diagnoses in 3q29Del cases to population prevalence values, one-sample proportion tests with Yates’ continuity correction were implemented using the stats R package [30]. To compare sex distribution between 3q29Del participants and controls, Pearson’s chi square test was implemented using the stats R package [30]. To compare age distribution in 3q29Del participants and controls, two sample t-test was implemented using the stats R package [30]. To compare scores in 3q29Del participants to mean values for children with idiopathic ASD, one sample t-test was implemented using the stats R package [30]. Odds ratios and p values were calculated using the fmsb R package [32]. Figures were generated using the plotly and ggplot2 R packages [33][34].

#### Sensitivity Analysis

The questionnaires for 90 participants with 3q29Del (96.8%) were completed by a parent or guardian (“parent-registered”), while 3 participants with 3q29Del (3.2%) completed all questionnaires themselves (“self-registered”). All control participants were parent-registered. To assess whether responses from the self-registered 3q29Del participants were influencing the results, self-registrants were removed and the data were re-analyzed. Self-registrants were not found to have a significant effect on the analyses (Tables S2 and S3). All results include both parent- and self-registrants.

## RESULTS

### Self-report of neuropsychiatric diagnosis in 3q29Del

Self-report of neuropsychiatric diagnoses in our 3q29Del study subjects (Table 2) revealed a higher prevalence of neuropsychiatric disorder diagnoses compared to controls, including anxiety (28.0%), and compared to general population frequencies, including ASD (29.0%, Figure 2A) and GDD/MR (59.1%) (Table 2), confirming prior work by our group [5]. Reported rates of conduct disorder (1.1% vs. 3.5%) and oppositional defiant disorder (3.2% vs. 3.5%) were similar to those observed in the general population. While a small proportion of participants reported diagnoses of bipolar/manic depression (4.3%), depression (6.5%), and schizophrenia (4.3%), we focused on ASD due to the young age (mean = 10.0 years) of our study population, since many study participants have not reached the age of risk for schizophrenia and other adult-onset disorders. Despite this young age, the self-reported rate of SZ diagnoses in our adult study subjects (age > 18 years, n = 13) was 15-30 times higher than expected (15.4% compared to an expected 0.5-1% in the general population; n = 2) [35–39] and the frequency of bipolar disorder was ∼1.8 times higher than expected [40]. A summary of neuropsychiatric diagnoses can be found in Table 2.

**Table 2:**
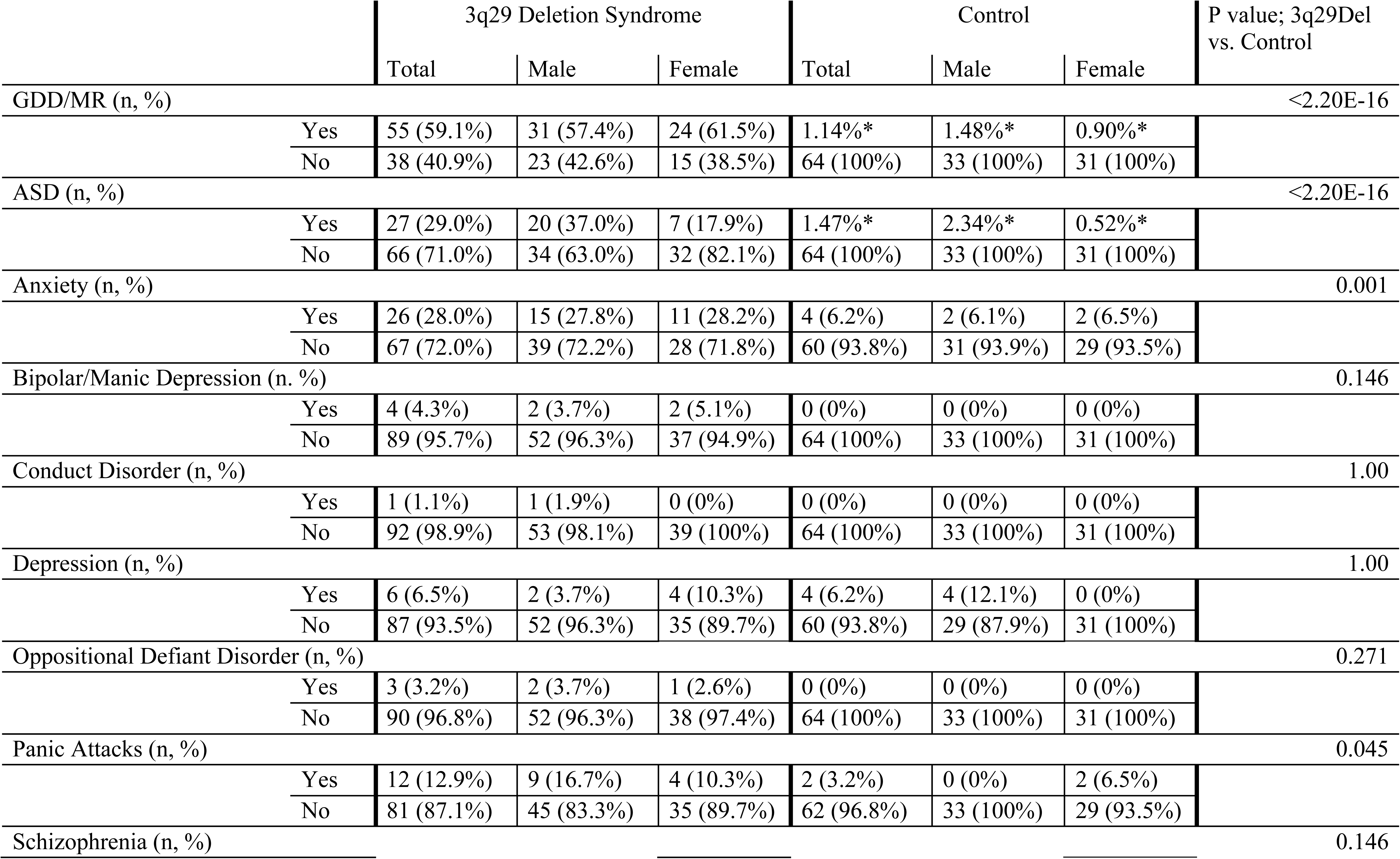
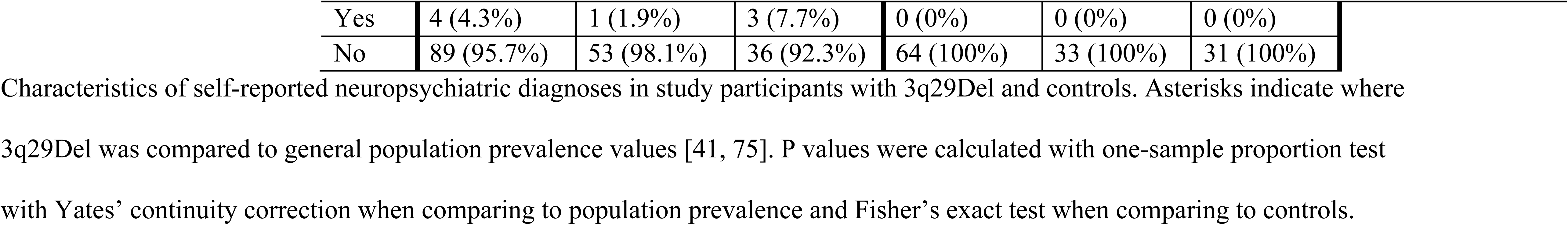
Self-reported neuropsychiatric diagnoses.

|  | 3q29 Deletion Syndrome |  |  | Control |  |  | P value; 3q29Del vs. Control |
| --- | --- | --- | --- | --- | --- | --- | --- |
|  | Total | Male | Female | Total | Male | Female |  |
| GDD/MR (n, %) |  |  |  |  |  |  | <2.20E-16 |
| Yes | 55 (59.1%) | 31 (57.4%) | 24 (61.5%) | 1.14%* | 1.48%* | 0.90%* |  |
| No | 38 (40.9%) | 23 (42.6%) | 15 (38.5%) | 64 (100%) | 33 (100%) | 31 (100%) |  |
| ASD (n, %) |  |  |  |  |  |  | <2.20E-16 |
| Yes | 27 (29.0%) | 20 (37.0%) | 7 (17.9%) | 1.47%* | 2.34%* | 0.52%* |  |
| No | 66 (71.0%) | 34 (63.0%) | 32 (82.1%) | 64 (100%) | 33 (100%) | 31 (100%) |  |
| Anxiety (n, %) |  |  |  |  |  |  | 0.001 |
| Yes | 26 (28.0%) | 15 (27.8%) | 11 (28.2%) | 4 (6.2%) | 2 (6.1%) | 2 (6.5%) |  |
| No | 67 (72.0%) | 39 (72.2%) | 28 (71.8%) | 60 (93.8%) | 31 (93.9%) | 29 (93.5%) |  |
| Bipolar/Manic Depression (n, %) |  |  |  |  |  |  | 0.146 |
| Yes | 4 (4.3%) | 2 (3.7%) | 2 (5.1%) | 0 (0%) | 0 (0%) | 0 (0%) |  |
| No | 89 (95.7%) | 52 (96.3%) | 37 (94.9%) | 64 (100%) | 33 (100%) | 31 (100%) |  |
| Conduct Disorder (n, %) |  |  |  |  |  |  | 1.00 |
| Yes | 1 (1.1%) | 1 (1.9%) | 0 (0%) | 0 (0%) | 0 (0%) | 0 (0%) |  |
| No | 92 (98.9%) | 53 (98.1%) | 39 (100%) | 64 (100%) | 33 (100%) | 31 (100%) |  |
| Depression (n, %) |  |  |  |  |  |  | 1.00 |
| Yes | 6 (6.5%) | 2 (3.7%) | 4 (10.3%) | 4 (6.2%) | 4 (12.1%) | 0 (0%) |  |
| No | 87 (93.5%) | 52 (96.3%) | 35 (89.7%) | 60 (93.8%) | 29 (87.9%) | 31 (100%) |  |
| Oppositional Defiant Disorder (n, %) |  |  |  |  |  |  | 0.271 |
| Yes | 3 (3.2%) | 2 (3.7%) | 1 (2.6%) | 0 (0%) | 0 (0%) | 0 (0%) |  |
| No | 90 (96.8%) | 52 (96.3%) | 38 (97.4%) | 64 (100%) | 33 (100%) | 31 (100%) |  |
| Panic Attacks (n, %) |  |  |  |  |  |  | 0.045 |
| Yes | 12 (12.9%) | 9 (16.7%) | 4 (10.3%) | 2 (3.2%) | 0 (0%) | 2 (6.5%) |  |
| No | 81 (87.1%) | 45 (83.3%) | 35 (89.7%) | 62 (96.8%) | 33 (100%) | 29 (93.5%) |  |
| Schizophrenia (n, %) |  |  |  |  |  |  | 0.146 |

|  |  |  |  |  |  |  |
| --- | --- | --- | --- | --- | --- | --- |
| Yes | 4 (4.3%) | 1 (1.9%) | 3 (7.7%) | 0 (0%) | 0 (0%) | 0 (0%) |
| No | 89 (95.7%) | 53 (98.1%) | 36 (92.3%) | 64 (100%) | 33 (100%) | 31 (100%) |

Characteristics of self-reported neuropsychiatric diagnoses in study participants with 3q29Del and controls. Asterisks indicate where 3q29Del was compared to general population prevalence values [41, 75]. P values were calculated with one-sample proportion test with Yates' continuity correction when comparing to population prevalence and Fisher's exact test when comparing to controls.

### SRS, SCQ, ASSQ, and CBCL/ABCL scores

In 3q29 deletion study subjects, the mean SRS score was in the moderate range (*T-score* = 71.8), the mean ASSQ score was in the clinical range (mean = 22.2), and the mean CBCL/ABCL score was in the borderline range (*T-score* = 62.5). The mean SCQ score in 3q29 deletion carriers was at the extremely high end of the normal range (mean = 13.9, clinical cutoff = 15) and elevated as compared to controls (mean = 3.5). Mean scores for typically developing controls were all in the normal range (SRS *T-score* = 45.9, ASSQ mean = 2.2, CBCL/ABCL *T-score* = 41.8, SCQ mean = 3.5) (Figure 1). Participants with 3q29Del scored significantly higher than typically developing controls on all four scales (p < 3.0E-12, Table S4).

**Figure 1.**
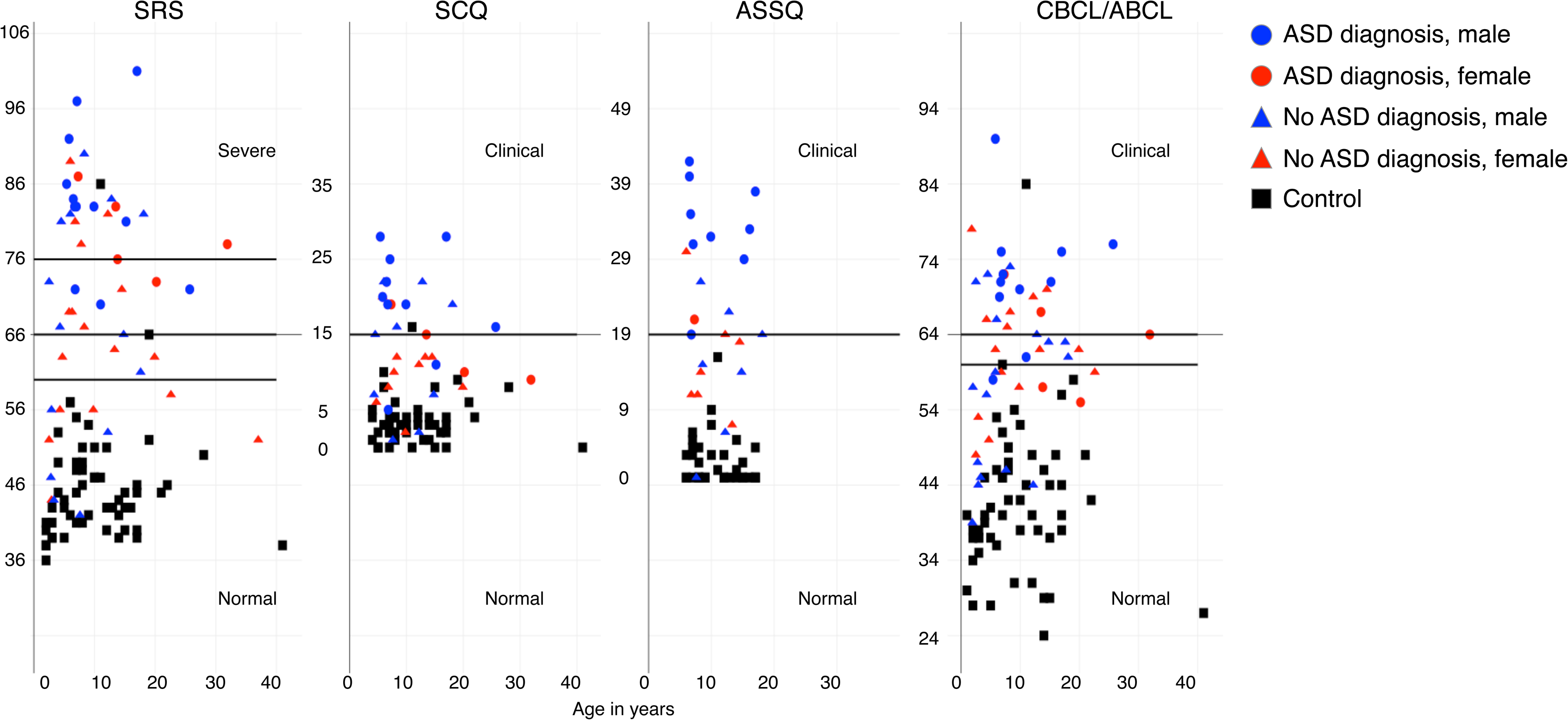
Score distribution for 3q29Del and controls on the SRS, SCQ, ASSQ, and CBCL/ABCL. Total scores on the SRS (n=48 3q29Del, 56 control), SCQ (n=33 3q29Del, 46 control), ASSQ (n=24 3q29Del, 35 control), and CBCL/ABCL (n=48 3q29Del, 57 control) for registry participants. Self-reported diagnosis of ASD is denoted by shape (circle/ASD, triangle/no ASD), and sex of participant is denoted by color (red/female, blue/male). Controls are shown in black.

### Standardized scores stratified by ASD diagnosis

Next, we examined the relationship between SRS scores and reported ASD diagnosis, to determine whether the score inflation we observed in our study population as a whole was largely due to the increased prevalence of ASD. As expected, we observed that individuals with 3q29Del and an ASD diagnosis scored significantly higher than both controls and individuals with 3q29Del without an ASD diagnosis (3q29Del with ASD n = 17, *T-score* = 82.41; 3q29Del without ASD n = 31, *T-score* = 65.90; control n = 56, *T-score* = 45.90; p < 3.0E-13; Figure 2B).

**Figure 2.**
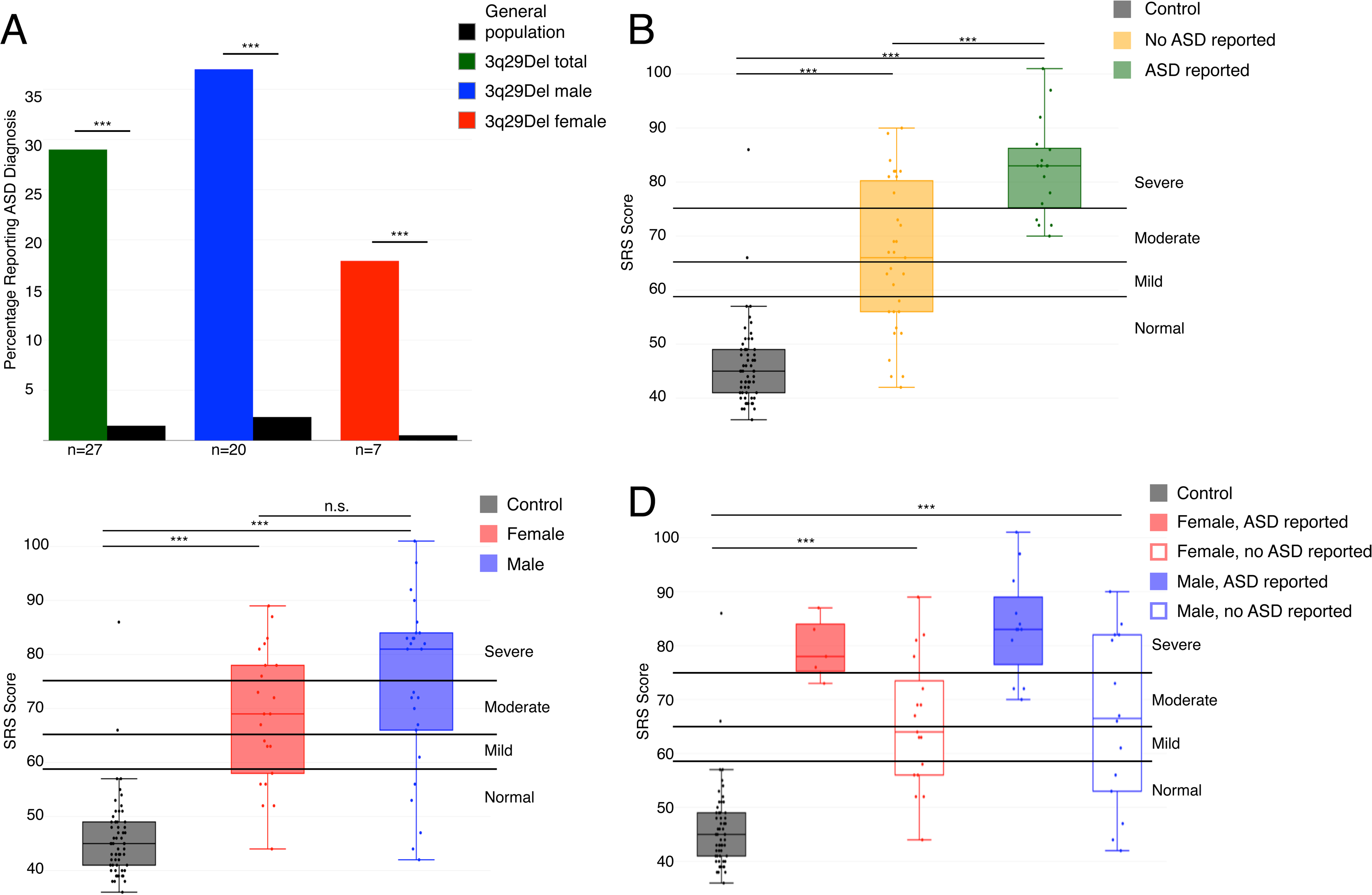
Comparison of ASD prevalence and SRS scores between 3q29Del and controls. **A)** Proportion of participants with 3q29Del self-reporting a diagnosis of ASD (not all respondents completed symptom questionnaires); 27 cases report an ASD diagnosis (green), comprised of 20 males (blue) and 7 females (red). Compared to general population frequencies (black), cases report significantly higher incidence of ASD. **B)** SRS scores split by control (n=59), 3q29Del not reporting an ASD diagnosis (n=31), and 3q29Del reporting an ASD diagnosis (n=17), showing a significant association between self-reported diagnostic status and SRS score. **C)** SRS scores split by sex, with control (n=59), 3q29Del female (n=22), and 3q29Del male (n=26), showing a lack of sex bias in scores for 3q29Del participants. **D)** SRS scores split by sex and self-reported diagnostic status, with control (n=59), 3q29Del female reporting ASD (n=5), 3q29Del female not reporting ASD (n=17), 3q29Del male reporting ASD (n=12), and 3q29Del male not reporting ASD (n=14), showing inflated scores for 3q29Del participants irrespective of sex or diagnostic status. ***, p<0.001

We were interested to observe that individuals with 3q29Del without an ASD diagnosis also scored significantly higher than controls (3q29Del without ASD n = 31, *T-score* = 65.90; control n = 56, *T-score* = 45.90; p = 2.16E-13; Figure 2B), indicating that increased SRS scores in individuals with 3q29Del are not driven by ASD diagnostic status alone (Table S5). Similar features were observed in the contribution of ASD diagnosis status to SCQ scores (Figure S1, Table S6).

### Standardized scores stratified by sex

Both males and females with 3q29Del reported a significantly increased frequency of ASD diagnoses, with a substantially greater burden for ASD on females with 3q29Del. Males with 3q29Del are at 16-fold increased risk for ASD as compared to the general population (37.0% vs. 2.34%, OR = 24.6, p = 6.06E-09) and females are at 34-fold risk compared to the general population (17.9% vs. 0.52%, OR = 41.8, p = 4.78E-05) (figure 2A) [41], resulting in a male:female ratio in our study population of 2:1, as compared to the general population ratio of 4:1. Taken together, this indicates that the 3q29 deletion elevates the risk for ASD in females more substantially than in males.

Based on the sex differences in ASD risk for individuals with 3q29Del, we also examined possible sex differences in scores. We found that both males and females with 3q29Del scored significantly higher than controls (3q29Del male n = 26, *T-score* = 74.31; control male n = 30, *T-score* = 45.80; p = 7.70E-11; 3q29Del female n = 22, *T-score* = 68.73; control female n = 26, *T-score* = 46.04; p = 7.42E-09); while 3q29Del males have higher scores than females, the differences are not statistically significant (3q29Del male n = 26, *T-score* = 74.31; 3q29Del female n = 22, *T-score* = 68.73; p > 0.05; Figure 2C). After stratifying our study population further by sex and ASD diagnosis status, we determined that both male and female 3q29Del participants without an ASD diagnosis had significantly higher scores than controls (3q29Del male without ASD n = 14, *T-score* = 66.29; control male n = 30, *T-score* = 45.80; p = 1.20E-06; 3q29Del female without ASD n = 17, *T-score* = 65.69; control female n = 26, *T-score* = 46.04; p = 5.04E-07; Figure 2D). Taken together, this suggests that increased SRS scores in individuals with 3q29Del are not driven by sex alone or by sex and ASD diagnosis status in combination (Table S5); rather, the presence of the deletion itself confers a greater risk for social disability. Furthermore, these data show an enrichment for female ASD in our study population, based on the reduction in male bias and the highly similar scores between males and females with 3q29Del, irrespective of ASD diagnosis status. Similar features were observed in the contribution of sex to SCQ scores (Figure S1, Table S6).

### ASD presentation of 3q29Del

While total scores on the SRS, SCQ, ASSQ, and CBCL/ABCL can give an indication of the overall level of impairment of individuals, sub-scores can reveal nuanced deficits in specific behavioral domains. To this end, we analyzed all SRS sub-scales (Social Awareness, Social Cognition, Social Communication, Social Motivation, Restricted Interests and Repetitive Behaviors, and Social Communication and Interaction) to better understand the extent of social disability in our study population; our goal was to determine whether our observed total score inflation was due to a specific severe deficit in a few domains, or if individuals with 3q29Del showed high scores across all sub-scales. The mean score for the Restricted Interests and Repetitive Behaviors sub-scale was in the severe range (*T-score* = 77.3). Mean scores for Social Awareness (*T-score* = 67.3), Social Cognition (*T-score* = 69.1), Social Communication (*T-score* = 69.7), and Social Communication and Interaction (*T-score* = 69.5) were all in the moderate range. Notably, the mean score for Social Motivation was in the mild range (*T-score* = 62.1, Figure 3A, Table 3). This sub-score profile is strikingly different from that reported in studies of idiopathic ASD, where children tend to score equally high on all sub-scales (3q29Del Social Motivation *T-score* = 62.1, idiopathic ASD Social Motivation *T-score* = 78.4, p = 7.66E-11) [42]. This atypical behavioral profile is supported by clinical data; direct assessment of individuals with 3q29Del by clinicians affiliated with the Emory 3q29 Project (http://genome.emory.edu/3q29/, [43]) show less impaired social motivation as compared to children with idiopathic ASD.

**Figure 3.**
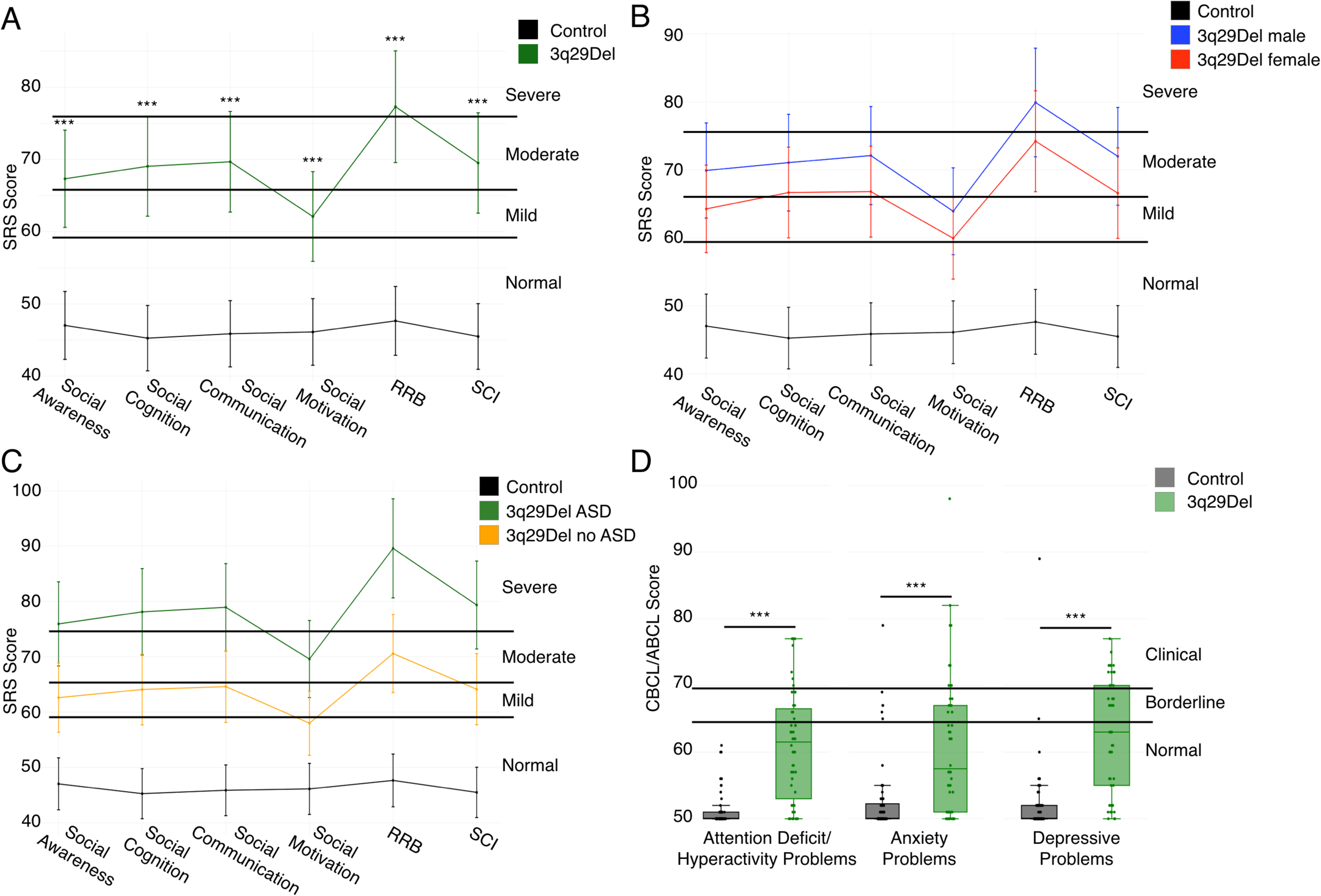
Comparison of SRS sub-scales and CBCL/ABCL DSM-oriented sub-scales between 3q29Del and controls. **A)** Profile of individuals with 3q29Del (n=48) and controls (n=59) across SRS sub-scales, showing moderate to severe impairment of 3q29Del participants in all domains except Social Motivation (RRB, Restricted Interests and Repetitive Behaviors; SCI, Social Communication and Interaction). **B)** Profile of 3q29Del males (n=26) and females (n=22) and controls (n=59) across SRS sub-scales, showing that 3q29Del males and females both score significantly higher than controls and that there are no significant differences in score between males and females. **C)** Profile of 3q29Del participants reporting an ASD diagnosis (n=17) and participants not reporting an ASD diagnosis (n=31) and controls (n=59) across SRS sub-scales, showing that 3q29Del participants score significantly higher than controls irrespective of ASD status, with 3q29Del participants reporting an ASD diagnosis scoring significantly higher than those not reporting an ASD diagnosis. **D)** Profile of 3q29Del participants (n=48) and controls (n=57) across 3 DSM-oriented sub-scales from the CBCL and ABCL, showing significantly increased pathology in 3q29Del participants in all 3 domains. ***, p<0.001

**Table 3:** SRS sub-scale score comparison stratified by genotype, ASD status, and sex.

|  | Social Awareness |  | Social Cognition |  | Social Communication |  | Social Motivation |  | RRB |  | SCI |  |
| --- | --- | --- | --- | --- | --- | --- | --- | --- | --- | --- | --- | --- |
| | Mean ± SD | P value | Mean ± SD | P value | Mean ± SD | P value | Mean ± SD | P value | Mean ± SD | P value | Mean ± SD | P value |
| **Genotype** |  |  |  |  |  |  |  |  |  |  |  |  |
| Control | 47.04 ± 8.88 | - | 45.27 ± 7.64 | - | 45.88 ± 8.14 | - | 46.13 ± 7.66 | - | 47.66 ± 8.51 | - | 45.50 ± 7.74 | - |
| 3q29Del | 67.33 ± 13.28 | 1.45E-13 | 69.06 ± 15.51 | 1.20E-15 | 69.69 ± 13.92 | <2.00E-16 | 62.10 ± 13.52 | 1.62E-10 | 77.31 ± 14.25 | <2.00E-16 | 69.52 ± 14.63 | <2.00E-16 |
| **Sex** |  |  |  |  |  |  |  |  |  |  |  |  |
| Male control | 45.97 ± 9.15 | - | 45.23 ± 7.42 | - | 45.73 ± 6.34 | - | 46.57 ± 6.58 | - | 47.63 ± 5.59 | - | 45.37 ± 6.60 | - |
| Male 3q29Del | 69.92 ± 13.92 | 7.61E-09 | 71.08 ± 17.65 | 1.07E-08 | 72.12 ± 15.47 | 1.74E-10 | 63.92 ± 15.23 | 3.39E-06 | 79.92 ± 14.82 | 2.84E-13 | 72.00 ± 16.34 | 7.12E-10 |
| Female control | 48.27 ± 8.56 | - | 45.31 ± 8.03 | - | 46.04 ± 9.95 | - | 45.62 ± 8.85 | - | 47.69 ± 11.08 | - | 45.65 ± 9.01 | - |
| Female 3q29Del | 64.27 ± 12.08 | 4.52E-06 | 66.68 ± 12.51 | 1.29E-08 | 66.82 ± 11.52 | 2.99E-08 | 59.95 ± 11.14 | 1.36E-05 | 74.23 ± 13.22 | 1.30E-09 | 66.59 ± 12.02 | 1.87E-08 |
| **ASD Status** |  |  |  |  |  |  |  |  |  |  |  |  |
| Control | 47.04 ± 8.88 | - | 45.27 ± 7.64 | - | 45.88 ± 8.14 | - | 46.13 ± 7.66 | - | 47.66 ± 8.51 | - | 45.50 ± 7.74 | - |
| No ASD diagnosis 3q29Del | 62.61 ± 13.20 | 5.17E-09 | 64.10 ± 15.80 | 5.77E-11 | 64.61 ± 13.54 | 1.20E-12 | 58.00 ± 13.15 | 1.42E-06 | 70.58 ± 11.50 | <2.00E-16 | 64.13 ± 14.39 | 5.19E-12 |
| ASD diagnosis 3q29Del | 75.94 ± 8.33 | 2.43E-15 | 78.12 ± 10.18 | <2.00E-16 | 78.94 ± 9.18 | <2.00E-16 | 69.59 ± 10.99 | 5.12E-12 | 89.59 ± 10.04 | <2.00E-16 | 79.35 ± 9.02 | <2.00E-16 |

Comparison of mean scores on the SRS sub-scales between study participants with 3q29Del and controls. 3q29Del participants were stratified by ASD status and sex for further analysis. P values were calculated using simple linear regression, adjusting for age, race, and sex.

#### ASD presentation stratified by sex

To determine whether this unusual SRS sub-score profile was influenced by sex, we examined profiles of male and female 3q29 deletion carriers separately. We found that the shape of the profiles were identical, with males scoring on average 5 points higher than females on every sub-scale (n = 26 male, 22 female; p > 0.05; Figure 3B; Table 3), demonstrating that the social disability in 3q29Del is not qualitatively different between males and females.

#### ASD presentation stratified by ASD diagnosis

We then stratified our study subjects according to reported ASD diagnosis status and examined subscale scores separately for 3q29Del individuals reporting a diagnosis of ASD and those not reporting a diagnosis of ASD. We observed that the shape of the profile is shared between 3q29Del individuals reporting a diagnosis of ASD and those not reporting a diagnosis of ASD, with individuals reporting a diagnosis of ASD scoring on average 10-15 points higher on every sub-scale (Figure 3C). As expected, 3q29Del participants with ASD scored significantly higher on all sub-scales than 3q29Del participants without ASD (n = 17 with ASD, 31 without ASD; p < 0.005; Table 3); however, 3q29Del participants without ASD still scored significantly higher than controls on all sub-scales (n = 31 without ASD, 56 control; p < 5.0E-05; Table 3).

### Additional neuropsychiatric phenotypes in 3q29Del

To further assess behavioral features associated with the 3q29 deletion, we examined the DSM-oriented Attention Deficit/Hyperactivity Problems, Anxiety Problems, and Depressive Problems sub-scales from the CBCL and ABCL. These DSM-oriented sub-scales align with neuropsychiatric diagnoses reported by individuals with 3q29Del [5]. Individuals with 3q29Del scored significantly higher than typically developing controls on all three scales (3q29Del Attention Deficit/Hyperactivity Problems *T-score* = 61.0, control Attention Deficit/Hyperactivity Problems *T-score* = 51.3, 3q29Del Anxiety Problems *T-score* = 60.9, control Anxiety Problems *T-score* = 52.9, 3q29Del Depressive Problems *T-score* = 62.7, control Depressive Problems *T-score* = 52.3, all p < 0.001, Figure 3D, Table S7), supporting previous reports of increased risk for neuropsychiatric phenotypes associated with the 3q29 deletion [5].

### Confounding due to heart defects and/or ID-related phenotypes

A previous study of 3q29Del by our group showed that approximately 25% of individuals with 3q29Del reported a congenital heart defect [5]. Early hypoxic insult due to a heart defect has been hypothesized to contribute to later neuropsychiatric and neurodevelopmental outcomes [44–49]. To determine if the high frequency of heart defects in our study population was driving adverse neurodevelopmental outcomes within 3q29Del cases, we implemented generalized linear and cumulative link models to assess the relationship between congenital heart defects and clinical ASD diagnosis, GDD/MR diagnosis, and age at walking, which has been reported to be a suitable proxy for ID in the absence of available IQ and adaptive behavior measures [50].

Congenital heart defects were not associated with self-reported ASD or GDD/MR diagnoses or age at walking (p > 0.05, Table S8). Individuals with 3q29Del are also commonly diagnosed with mild to moderate ID [5]. To ask whether ASD phenotypes or ASD features were disproportionately overrepresented in individuals with more pronounced ID-related phenotypes and/or heart defects, we stratified the data according to these phenotypes. Within our 3q29Del study population, congenital heart defects were associated with significantly increased scores on the SCQ and CBCL/ABCL (p < 0.05); however, reported GDD/MR diagnosis and age at walking were not significantly associated with scores on the SRS, SCQ, ASSQ, or CBCL/ABCL (p > 0.05, Table S9). These data indicate that ID-related phenotypes were not driving the increased scores in our study population.

## DISCUSSION

Previous studies have found enrichment of the 3q29 deletion in large samples ascertained based on clinical ASD diagnosis [17, 18]. We have approached the association of 3q29Del with ASD from a different angle; by ascertaining subjects with 3q29Del and investigating the prevalence of reported ASD diagnosis and ASD-related phenotypes, the current study complements the existing literature, providing additional evidence for the 3q29 deletion as a genetic risk factor for ASD. Notably, the male:female ratio of self-reported ASD diagnosis in our study population is 2:1. This is a reduction from the 4:1 male bias observed in idiopathic ASD in the general population. A substantial reduction in male bias in ASD prevalence has been observed in studies of other CNVs and single-gene mutations; a recent study has shown that as the severity of a mutation increases, the sex ratio in ASD prevalence approaches 1:1 [51]. Taken together, this suggests that the 3q29 deletion is approaching the severe end of the spectrum of ASD-associated mutations.

We have shown that compared to typically developing children, our 3q29Del sample is significantly enriched for ASD features and other behavioral problems, irrespective of a clinical ASD diagnosis. This finding is particularly concerning; while individuals with 3q29Del who have an ASD diagnosis tend to score higher on symptomology scales overall, 3q29Del individuals without an ASD diagnosis still score significantly higher than typically developing children. This indicates several possible explanations: a) an enrichment for ASD features or social disability that falls short of diagnostic criteria, b) possible undiagnosed ASD in our study population, or c) non-specificity of the SRS, and potentially SCQ, for phenotypes other than ASD, such as anxiety. The possibility of undiagnosed ASD in our study population is aligned with anecdotal reports from parents of our study participants, where they have reported concerns about atypical social development that do not appear to have been addressed using gold-standard ASD evaluations. Based on the elevated symptomology scores in our study population, the substantially increased risk for ASD associated with the 3q29 deletion, and the apparent severity of the 3q29 deletion, our data suggest that gold-standard ASD evaluations should be the recommended standard of care for individuals diagnosed with 3q29Del. If implemented, this practice would enable patients to gain access to early interventions, treatments, and therapeutic programs that are known to improve later outcomes.

Based on the SRS sub-scales, participants with 3q29Del display a strikingly different behavioral profile as compared to a study of children with idiopathic ASD [42]. Male and female 3q29Del individuals show substantially less impaired social motivation in the context of an otherwise typical ASD profile, with the most severe deficits in the Restricted Interests and Repetitive Behaviors domain. This profile is also observed when dividing scores for 3q29Del participants based on reported ASD diagnosis. This qualitative difference from idiopathic ASD may serve as an inroad to therapeutic interventions in 3q29Del, as well as an investigative inroad to a distinct subtype of ASD. Because social motivation appears to be relatively well-preserved in 3q29Del, this suggests that therapies such as cognitive-behavioral therapy to teach social skills and effective strategies for social interaction may be particularly successful in this patient population.

Some facets of the difference in ASD features between 3q29Del and idiopathic ASD are recapitulated by the scores on the Withdrawn sub-scale of the CBCL and ABCL. Previous studies utilizing the CBCL in idiopathic ASD have found that mean scores for participants with ASD are in the borderline range, with over 50% of subjects scoring in the borderline or clinical range [52, 53]. While 3q29Del participants generally, as well as males and females separately, score significantly higher than controls, their mean score is still in the normal range (Figure S2A and B). However, 60% of 3q29Del participants reporting an ASD diagnosis score in the borderline or clinical range (Figure S2C, Table S10), which is in line with what is expected based on studies of idiopathic ASD [52, 53]. This is in conflict with the relatively well-preserved social motivation in 3q29Del individuals with ASD identified in our analysis of the SRS sub-scales and suggests that a more refined analysis is merited to identify the true degree of social disability in this population.

We tested the hypothesis that the score inflation observed in our 3q29Del study subjects may be due to the high prevalence of developmental delay or congenital heart defects [5]. Our available data do not support this hypothesis, and instead reveal that social disability is equally distributed in our study population. Lack of direct measures of intellectual disability, and errors or missing data in self-report measures, may obscure this relationship; however, numerous studies of the relationship between ID and ASD in genomic disorders suggests that when the population is stratified by the presence of a specific genetic variant, the association between these two phenotypes diminishes. A large study of several genetic disorders showed that the prediction of genetic diagnosis based on ADI-R scores was not confounded by IQ [54]; a study of 7q11.23 duplication found that IQ was not significantly associated with ASD status [55]; and multiple studies of 22q11.2 deletion have shown that IQ is not significantly associated with SRS score, ASD severity, and ASD status [56–58]. A question ripe for future investigation is the potential role for microcephaly in the ASD-related phenotypes observed in 3q29Del.

Microcephaly, ASD, and ID are associated with the 16p11.2 duplication [21]; microcephaly has been shown to be associated with ASD and ID in probands with pathogenic CNVs [59]; and children with “complex autism”, defined as ASD with microcephaly and/or dysmorphology, have significantly lower cognitive function than children with “essential autism” [60]. Reports have shown a high prevalence of microcephaly in 3q29Del [3, 4, 12]; however, this question was not probed in the current study due to the high rate (>50%) of 3q29Del participants responding “Unsure” to the medical history questionnaire regarding their child’s head circumference at birth, rendering this data unreliable. Ongoing studies with direct evaluation of study subjects [43] will address these questions.

While this study is the most comprehensive study of behavioral phenotypes in 3q29Del to date, it is not without limitations. All of the data used in the present study were collected from questionnaires completed by the parents and guardians of individuals with 3q29Del, which introduces several potential sources of bias. Some studies have questioned the validity and reliability of parent-report data [61]; however, a recent study in Williams syndrome patients has shown that parents are more accurate in predicting their child’s social behaviors than the child themselves [62]. The responses to the medical and demographic questionnaire are more likely to include error due to the fact that the data is retrospective. By limiting our study to only a few key points in the medical history (heart defects, age at walking, and ID/ASD diagnosis) we aimed to reduce recall errors; however, we only had proxies for ID, rather than direct evaluation of cognitive ability. Further, the sample sizes for our stratified analyses were small, rendering them underpowered; while the differences between males and females were not statistically significant, males do score higher than females on all measures. Studies with larger sample size will be better able to assess the importance of and estimate the true effect size of any difference between males and females. Additionally, there is likely ascertainment bias within our sample. First, our sample of 93 individuals with 3q29Del is 87.1% white, indicating that we are not adequately reaching minority populations. Second, parents that register their children and complete in-depth questionnaires are likely to be highly motivated, possibly because their children experience significant morbidity – a potential indication that we are sampling from the extreme of the phenotypic distribution of 3q29Del. Thus, scores on the standardized questionnaires, as well as rates of heart defects and clinical neuropsychiatric diagnoses, may be higher in our study sample than in the general 3q29Del population. Additionally, the odds ratios calculated for the increased risk for ASD associated with the 3q29 deletion may also be overestimated, due to the combined effects of self-report data and ascertainment bias; however, if this increased risk is replicated using gold-standard diagnostic measures, it could provide valuable insight into possible sex-specific effects of the deletion. Finally, the lack of observed association between congenital heart defects and neurodevelopmental outcomes may be obscured by the high rate of patent ductus arteriosus in 3q29 deletion syndrome [5], which is a relatively mild heart defect; however, the low number of participants with different types of heart defects rendered analyses to assess their associations with neurodevelopment underpowered (Table S11). Ongoing studies by the Emory 3q29 Project (http://genome.emory.edu/3q29/), including direct in-person patient evaluations [43] aim to address some of the weaknesses of the present work by performing comprehensive gold-standard evaluations by expert clinicians.

While direct in-person evaluations are the ideal method to corroborate the findings of this study, the low population frequency of the 3q29 deletion and geographic dispersal of our study population (Figure S3) renders this approach infeasible for a large number of study subjects. However, a small number of 3q29 deletion study subjects have been directly assessed as part of the Emory 3q29 Project (http://genome.emory.edu/3q29/). We confirm high concordance between registry-leveraged data and gold-standard direct evaluation, as all participants qualifying for an ASD diagnosis based on gold-standard evaluation have clinically significant scores on the SRS and all participants reporting an ASD diagnosis qualified for an ASD diagnosis after gold-standard assessment by the Emory 3q29 Project team (Table S12). Notably, one participant that did not report a prior diagnosis of ASD received an ASD diagnosis after assessment by our team, supporting our hypothesis that ASD may be underdiagnosed in the 3q29Del population. Five additional participants with a clinically significant SRS score did not qualify for an ASD diagnosis, suggesting that the SRS is not selectively identifying children with ASD in participants with 3q29Del, possibly due to the high rates of reported anxiety in our study population. However, this comparison does suggest that our analysis, though based on self-report data, reveals valid conclusions about behavioral phenotypes in 3q29 deletion syndrome. For genetic syndromes with low population frequencies, data collection through remote means such as online patient registries remains a valuable phenotyping tool.

While the current understanding of the 3q29 deletion is still evolving, there are more well-understood CNV disorders that can be used as a comparison point to determine whether the social disability phenotypes described in this study are distinct to 3q29Del. These include Williams Syndrome (WS, or the 7q11.23 deletion), the reciprocal 7q11.23 duplication, 16p11.2 deletion and duplication, Smith-Magenis Syndrome (SMS), and 22q11.2 deletion. WS is typically associated with hyper-sociability [63], and patients with WS show more problems with social cognition than with pro-social behaviors [64], similar to what we have observed in our population of individuals with 3q29Del. However, the prevalence of restricted interests and repetitive behaviors appears to be lower in WS as compared to 3q29Del [64], and the mean SRS sub-scale Social Motivation score indicates enhanced social motivation in WS as compared to 3q29Del (WS mean *T-score* = 55.24, 3q29Del mean *T-score* = 62.1, p = 0.0005) [65]. Studies of the reciprocal 7q11.23 duplication showed that parent-reported ASD symptomology via standardized questionnaires was higher than ASD features as assessed by gold-standard instruments; that some probands had been diagnosed with ASD based on delayed speech and social anxiety but did not qualify for ASD via gold-standard measures; that substantially more males than females qualified for an ASD diagnosis; and that 7q11.23 duplication probands were indistinguishable from children with idiopathic ASD on measures of ASD severity and diagnosis status [55, 66, 67]. This is qualitatively different from our 3q29Del population; all of the participants with a prior ASD diagnosis who were later assessed by the Emory 3q29 Project team had their diagnosis confirmed using gold-standard measures (Table S12), the male:female ratio in our sample is 2:1, and we see significant differences between 3q29Del cases and idiopathic ASD [42] on the SRS Social Motivation sub-scale.

Similar to 7q11.23 duplication, ASD probands with 16p11.2 deletion or duplication were indistinguishable from idiopathic ASD probands [67]; probands with 16p11.2 deletion also have a significantly higher mean SRS score as compared to 3q29Del (16p11.2 mean *T-score* = 77.8, 3q29Del mean *T-score* = 71.8, p = 0.003) [22], and males with 16p11.2 deletion are at increased risk for ASD compared to females and are overrepresented when cases are ascertained based on neurodevelopmental disorders [68, 69], indicating a different sex-based ASD risk as compared to 3q29Del. A study of 16p11.2 duplication probands found that scores on the SRS Social Motivation sub-scale were not significantly different from controls and that ASD cases had specific impairments in social cognition and communication [70]; 3q29Del cases score significantly higher than controls on the SRS Social Motivation sub-scale, and do not have substantially higher scores on the Social Cognition or Social Communication sub-scales relative to the other SRS sub-scales.

A recent study of SMS showed that female probands scored higher than males on SRS sub-scales and the sex ratio of ASD was reversed, with more females than males qualifying for a diagnosis [71], which we do not observe in our 3q29Del study population. Finally, studies of 22q11.2 deletion show some similarities with 3q29Del, including SRS total scores that are not significantly different, high levels of ASD features in the absence of ASD diagnosis, and a male:female ASD ratio of approximately 1:1 [19, 57, 58, 72]; however, 22q11.2 deletion probands have a significantly lower mean ASSQ score as compared to 3q29Del (22q11.2 mean = 11, 3q29Del mean = 22.2, p = 0.00004), and 3q29Del cases have significantly higher scores on several CBCL/ABCL sub-scales (Table S13) [73, 74]. Taken together, this evidence suggests that while the ASD features in 3q29Del reported in this study share some characteristics with other CNV disorders, the complete constellation of symptoms is discrete from previously described genomic syndromes.

There are significant strengths of this study as compared to previous studies of 3q29Del. First, this is the largest cohort of individuals with 3q29Del ever assembled. This is a critical step in capturing the true phenotypic spectrum associated with the 3q29 deletion. Our use of standardized questionnaires allowed for comparison between ASD features present in 3q29Del and those reported in idiopathic ASD and ASD in other CNV disorders. Additionally, our online patient registry allows for remote data collection, which has enabled us to expand our sample size. This study has shown that high-quality, comprehensive medical history and symptomology data can be collected through an online patient registry, effectively reducing the patient-ascertainment burden associated with studying rare disorders. Taken together, these attributes make the present study an excellent complement to previously published case reports on individuals with 3q29Del; by capturing a larger patient base with systematic assessments, we are able to more accurately measure the presence of a variety of neuropsychiatric and neurodevelopmental phenotypes associated with the 3q29 deletion. The findings reported here indicate that comprehensive neuropsychiatric and neurodevelopmental assessments with gold standard tools are merited for individuals diagnosed with 3q29Del, and that such assessments should be the standard of care for this patient population.

## CONCLUSIONS

The present study confirms previous reports of phenotypes in 3q29Del, as well as expanding the spectrum of behavioral phenotypes associated with the deletion. We found that individuals with 3q29Del report a significantly higher prevalence of ASD diagnosis than the general population, and significantly elevated scores on the SRS, SCQ, ASSQ, and CBCL/ABCL irrespective of ASD diagnosis indicate significant social disability overall in our study population. Further, 3q29Del participants showed a distinct profile of ASD-related phenotypes on the SRS sub-scales, marked by less impaired scores on the Social Motivation sub-scale and extremely high scores on the Restricted Interests and Repetitive Behaviors sub-scale. This score profile is consistent between 3q29Del males and females and between 3q29Del participants with and without ASD, suggesting that it may be a hallmark behavioral feature of the syndrome and providing a potential therapeutic inroad for the treatment of individuals with 3q29Del. Finally, we identify a high degree of social disability in female 3q29Del participants; the 3q29 deletion elevates the risk ASD in females (OR=41.8, p=4.78E-05) more substantially than in males (OR=24.6, p=6.06E-09). These results demonstrate that there is a benefit to studying rare CNVs such as 3q29Del; studying a single genomic variant with large effect allows us to control for genetic etiology and unmask the mechanisms underlying the development of neuropsychiatric and neurodevelopmental disorders.

## Abbreviations

ASD: autism spectrum disorder
3q29Del: 3q29 deletion syndrome
SRS: Social Responsiveness Scale
SCQ: Social Communication Questionnaire
ASSQ: Autism Spectrum Screening Questionnaire
CBCL: Child Behavior Checklist
ABCL: Adult Behavior Checklist
DSM: Diagnostic and Statistical Manual
SZ: schizophrenia
ADHD: attention deficit/hyperactivity disorder
CNV: copy number variation
ID: intellectual disability
GDD: global developmental delay
MR: mental retardation
WS: Williams Syndrome
SMS: Smith-Magenis Syndrome

## DECLARATIONS

### Ethics approval and consent to participate

This study was approved by Emory University’s Institutional Review Board (IRB00064133). All study subjects gave informed consent prior to participating in this study.

### Consent for publication

Not applicable.

### Availability of data and material

The datasets used and analyzed during the current study are available from the corresponding author on reasonable request.

### Authors’ contributions

RMP performed the statistical analysis, produced all figures and tables, and wrote the manuscript. MMM collected the data. MPE helped with statistical analyses and interpretation. MMM, CK, and CAS helped with data interpretation. MEZ and JGM edited the manuscript and provided guidance on analyzing and interpreting data. JGM was the principle investigator responsible for study direction. All authors participated in commenting on the drafts and have read and approved the final manuscript.

## Acknowledgements

We gratefully acknowledge our study population, the 3q29 deletion community, for their participation and commitment to research. This manuscript is currently available on the bioRxiv preprint server (https://doi.org/10.1101/386243). We also acknowledge the contributions of the members of the Emory 3q29 Project: Hallie Averbach, Emily Black, Gary J Bassell, T Lindsey Burrell, Grace Carlock, Tamara Caspary, Joseph F Cubells, David Cutler, Paul A Dawson, Roberto Espana, Michael J Gambello, Katrina Goines, Henry R Johnston, Sookyong Koh, Elizabeth J Leslie, Longchuan Li, Bryan Mak, Tamika Malone, Trenell Mosley, Derek Novacek, Ryan Purcell, Timothy Rutkowski, Rossana Sanchez, Jason Schroeder, Esra Sefik, Brittney Sholar, Sarah Shultz, Nikisha Sisodiya, Steven Sloan, Elaine F Walker, Stephen T Warren, David Weinshenker, Zhexing Wen, and Mike Zinsmeister.

## Additional Files

File name: Supplemental Information

File format: Microsoft Word document (.docx) Title of data: Supplementary figures and tables

Description of data: Supplementary figures are (S1) SCQ scores split by ASD status, sex, and ASD status/sex, (S2) CBCL/ABCL Withdrawn sub-scale scores split by genotype, sex, and ASD status, and (S3) geographic distribution of study participants with 3q29Del. Supplementary tables are (S1) questionnaire demographics for the medical questionnaire, SRS, SCQ, ASSQ, and CBCL/ABCL; (S2 and S3) sensitivity analysis description and results for effect of 3q29Del self-registrants; (S4) comparison of scores on all four scales for 3q29Del versus control; (S5) SRS score comparison stratified by ASD status and sex; (S6) SCQ score comparison stratified by ASD status and sex; (S7) CBCL/ABCL DSM-oriented sub-scale score comparison; (S8) contribution of congenital heart defects to phenotypes of interest; (S9) test for confounding factors contributing to symptomology questionnaire scores; (S10) CBCL/ABCL Withdrawn sub-scale score comparison; (S11) heart defects present in study sample; (S12) comparison of 3q29 registry-leveraged and gold-standard phenotyping measures; and (S13) comparison of CBC/ABCL sub-scale scores between 3q29Del and 22q11.2 deletion.

## REFERENCES

1. Stefansson H, Meyer-Lindenberg A, Steinberg S, Magnusdottir B, Morgen K, Arnarsdottir S, Bjornsdottir G, Walters GB, Jonsdottir GA, Doyle OM, et al: CNVs conferring risk of autism or schizophrenia affect cognition in controls. Nature 2014, 505:361–366.

2. Kendall KM, Rees E, Escott-Price V, Einon M, Thomas R, Hewitt J, O’Donovan MC, Owen MJ, Walters JTR, Kirov G: Cognitive Performance Among Carriers of Pathogenic Copy Number Variants: Analysis of 152,000 UK Biobank Subjects. Biol Psychiatry 2017, 82:103–110.

3. Willatt L, Cox J, Barber J, Cabanas ED, Collins A, Donnai D, FitzPatrick DR, Maher E, Martin H, Parnau J, et al: 3q29 microdeletion syndrome: clinical and molecular characterization of a new syndrome. American Journal of Human Genetics 2005, 77:154–160.

4. Ballif BC, Theisen A, Coppinger J, Gowans GC, Hersh JH, Madan-Khetarpal S, Schmidt KR, Tervo R, Escobar LF, Friedrich CA, et al: Expanding the clinical phenotype of the 3q29 microdeletion syndrome and characterization of the reciprocal microduplication. Molecular Cytogenetics 2008, 1:8.

5. Glassford MR, Rosenfeld JA, Freedman AA, Zwick ME, Mulle JG, Unique Rare Chromosome Disorder Support G: Novel features of 3q29 deletion syndrome: Results from the 3q29 registry. American Journal of Medical Genetics Part A 2016, 170A:999–1006.

6. Cobb W, Anderson A, Turner C, Hoffman RD, Schonberg S, Levin SW: 1.3 Mb de novo deletion in chromosome band 3q29 associated with normal intelligence in a child. European Journal of Medical Genetics 2010, 53:415–418.

7. Mulle JG: The 3q29 deletion confers >40-fold increase in risk for schizophrenia. Molecular Psychiatry 2015, 20:1028–1029.

8. Mulle JG, Dodd AF, McGrath JA, Wolyniec PS, Mitchell AA, Shetty AC, Sobreira NL, Valle D, Rudd MK, Satten G, et al: Microdeletions of 3q29 confer high risk for schizophrenia. American Journal of Human Genetics 2010, 87:229–236.

9. Marshall CR, Howrigan DP, Merico D, Thiruvahindrapuram B, Wu W, Greer DS, Antaki D, Shetty A, Holmans PA, Pinto D, et al: Contribution of copy number variants to schizophrenia from a genome-wide study of 41,321 subjects. Nature Genetics 2017, 49:27–35.

10. Kirov G, Pocklington AJ, Holmans P, Ivanov D, Ikeda M, Ruderfer D, Moran J, Chambert K, Toncheva D, Georgieva L, et al: De novo CNV analysis implicates specific abnormalities of postsynaptic signalling complexes in the pathogenesis of schizophrenia. Mol Psychiatry 2012, 17:142–153.

11. Szatkiewicz JP, O’Dushlaine C, Chen G, Chambert K, Moran JL, Neale BM, Fromer M, Ruderfer D, Akterin S, Bergen SE, et al: Copy number variation in schizophrenia in Sweden. Molecular Psychiatry 2014, 19:762.

12. Cox DM, Butler MG: A clinical case report and literature review of the 3q29 microdeletion syndrome. Clinical dysmorphology 2015, 24:89–94.

13. Città S, Buono S, Greco D, Barone C, Alfei E, Bulgheroni S, Usilla A, Pantaleoni C, Romano C: 3q29 microdeletion syndrome: Cognitive and behavioral phenotype in four patients. American Journal of Medical Genetics Part A 2013, 161A:3018–3022.

14. Sagar A, Bishop JR, Tessman DC, Guter S, Martin CL, Cook EH: Co-occurrence of autism, childhood psychosis, and intellectual disability associated with a de novo 3q29 microdeletion. American Journal of Medical Genetics Part A 2013, 161A:845–849.

15. Quintero-Rivera F, Sharifi-Hannauer P, Martinez-Agosto JA: Autistic and psychiatric findings associated with the 3q29 microdeletion syndrome: case report and review. American Journal of Medical Genetics Part A 2010, 152A:2459–2467.

16. Biamino E, Di Gregorio E, Belligni EF, Keller R, Riberi E, Gandione M, Calcia A, Mancini C, Giorgio E, Cavalieri S, et al: A novel 3q29 deletion associated with autism, intellectual disability, psychiatric disorders, and obesity. American Journal of Medical Genetics Part B, Neuropsychiatric Genetics 2016, 171B:290–299.

17. Itsara A, Cooper GM, Baker C, Girirajan S, Li J, Absher D, Krauss RM, Myers RM, Ridker PM, Chasman DI, et al: Population analysis of large copy number variants and hotspots of human genetic disease. American Journal of Human Genetics 2009, 84:148–161.

18. Sanders SJ, He X, Willsey AJ, Ercan-Sencicek AG, Samocha KE, Cicek AE, Murtha MT, Bal VH, Bishop SL, Dong S, et al: Insights into Autism Spectrum Disorder Genomic Architecture and Biology from 71 Risk Loci. Neuron 2015, 87:1215–1233.

19. Schneider M, Debbané M, Bassett AS, Chow EWC, Fung WLA, van den Bree M, Owen M, Murphy KC, Niarchou M, Kates WR, et al: Psychiatric disorders from childhood to adulthood in 22q11.2 deletion syndrome: results from the International Consortium on Brain and Behavior in 22q11.2 Deletion Syndrome. The American Journal of Psychiatry 2014, 171:627–639.

20. McDonald-McGinn DM, Sullivan KE, Marino B, Philip N, Swillen A, Vorstman JA, Zackai EH, Emanuel BS, Vermeesch JR, Morrow BE, et al: 22q11.2 deletion syndrome. Nat Rev Dis Primers 2015, 1:15071.

21. D’Angelo D, Lebon S, Chen Q, Martin-Brevet S, Snyder LG, Hippolyte L, Hanson E, Maillard AM, Faucett WA, Mace A, et al: Defining the Effect of the 16p11.2 Duplication on Cognition, Behavior, and Medical Comorbidities. JAMA Psychiatry 2016, 73:20–30.

22. Hanson E, Bernier R, Porche K, Jackson FI, Goin-Kochel RP, Snyder LG, Snow AV, Wallace AS, Campe KL, Zhang Y, et al: The cognitive and behavioral phenotype of the 16p11.2 deletion in a clinically ascertained population. Biological Psychiatry 2015, 77:785–793.

23. Mervis CB, Klein-Tasman BP, Huffman MJ, Velleman SL, Pitts CH, Henderson DR, Woodruff-Borden J, Morris CA, Osborne LR: Children with 7q11.23 duplication syndrome: psychological characteristics. American Journal of Medical Genetics Part A 2015, 167:1436–1450.

24. Brunetti-Pierri N, Berg JS, Scaglia F, Belmont J, Bacino CA, Sahoo T, Lalani SR, Graham B, Lee B, Shinawi M, et al: Recurrent reciprocal 1q21.1 deletions and duplications associated with microcephaly or macrocephaly and developmental and behavioral abnormalities. Nature Genetics 2008, 40:1466–1471.

25. Constantino JN, Todd RD: The social responsiveness scale manual. 2 edn. Los Angeles: Western Psychological Services; 2012.

26. Rutter M, Bailey A, Lord C: The Social Communication Questionnaire - Manual. Los Angeles: Western Psychological Services; 2003.

27. Ehlers S, Gillberg C, Wing L: A screening questionnaire for Asperger syndrome and other high-functioning Autism Spectrum Disorders in school age children. Journal of Autism and Developmental Disorders 1999, 29.

28. Achenbach TM, Rescorla LA: Manual for the ASEBA School-Age Forms & Profiles. Burlington, VT: University of Vermont, Research Center for Children, Youth and Families; 2001.

29. Achenbach TM, Rescorla LA: Manual for the ASEBA Adult Forms & Profiles. Burlington, VT: University of Vermont, Research Center for Children, Youth and Families; 2003.

30. R Core Team: R: A language and environment for statistical computing. R Foundation for Statistical Computing, Vienna, Austria 2008.

31. Christensen RHB: ordinal - Regression Models for Ordinal Data. R package version 20156-28 2015.

32. Nakazawa M: fmsb: Functions for medical statistics book with some demographic data. R package version 063 2018.

33. Sievert C, Parmer C, Hocking T, Chamberlain S, Ram K, Corvellec M, Despouy P: plotly: Create Interactive Web Graphics via ‘plotly.js’. R package version 460 2017.

34. Wickham H: ggplot2: Elegant Graphics for Data Analysis. New York: Springer-Verlag; 2009.

35. Kessler RC, Birnbaum H, Demler O, Falloon IR, Gagnon E, Guyer M, Howes MJ, Kendler KS, Shi L, Walters E, Wu EQ: The prevalence and correlates of nonaffective psychosis in the National Comorbidity Survey Replication (NCS-R). Biol Psychiatry 2005, 58:668–676.

36. Wu EQ, Shi L, Birnbaum H, Hudson T, Kessler R: Annual prevalence of diagnosed schizophrenia in the USA: a claims data analysis approach. Psychol Med 2006, 36:1535–1540.

37. Desai PR, Lawson KA, Barner JC, Rascati KL: Estimating the direct and indirect costs for community-dwelling patients with schizophrenia. Journal of Pharmaceutical Health Services Research 2013, 4:187–194.

38. Saha S, Chant D, Welham J, McGrath J: A systematic review of the prevalence of schizophrenia. PLoS Med 2005, 2:e141.

39. Moreno-Kustner B, Martin C, Pastor L: Prevalence of psychotic disorders and its association with methodological issues. A systematic review and meta-analyses. PLoS One 2018, 13:e0195687.

40. Merikangas KR, He JP, Burstein M, Swanson SA, Avenevoli S, Cui L, Benjet C, Georgiades K, Swendsen J: Lifetime prevalence of mental disorders in U.S. adolescents: results from the National Comorbidity Survey Replication--Adolescent Supplement (NCS-A). J Am Acad Child Adolesc Psychiatry 2010, 49:980–989.

41. Christensen DL, Braun KVN, Baio J, Bilder D, Charles J, Constantino JN, Daniels J, Durkin MS, Fitzgerald RT, Kurzius-Spencer M, et al: Prevalence and Characteristics of Autism Spectrum Disorder Among Children Aged 8 Years - Autism and Developmental Disabilities Monitoring Network, 11 Sites, United States, 2012. MMWR Surveill Summ 2018, 65:1–23.

42. Torske T, Naerland T, Oie MG, Stenberg N, Andreassen OA: Metacognitive Aspects of Executive Function Are Highly Associated with Social Functioning on Parent-Rated Measures in Children with Autism Spectrum Disorder. Front Behav Neurosci 2017, 11:258.

43. Murphy MM, Lindsey Burrell T, Cubells JF, Espana RA, Gambello MJ, Goines KCB, Klaiman C, Li L, Novacek DM, Papetti A, et al: Study protocol for The Emory 3q29 Project: evaluation of neurodevelopmental, psychiatric, and medical symptoms in 3q29 deletion syndrome. BMC Psychiatry 2018, 18:183.

44. Rogers BT, Msali ME, Buck GM, Lyon NR, Norris MK, Roland JMA, Gingell RL, Cleveland DC, Pieroni DR: Neurodevelopmental outcome of infants with hypoplastic left heart syndrome. The Journal of Pediatrics 1995:496–498.

45. Forbess JM, Visconti KJ, Hancock-Friesen C, Howe RC, Bellinger DC, Jonas RA: Neurodevelopmental outcome after congenital heart surgery: Results from an institutional registry. Circulation 2002, 106:I-95-I-102.

46. Wernovsky G, Shillingford AJ, Gaynor JW: Central nervous system outcomes in children with complex congenital heart disease. Current Opinions in Cardiology 2005:94–99.

47. Karsdorp PA, Everaerd W, Kindt M, Mulder BJ: Psychological and cognitive functioning in children and adolescents with congenital heart disease: a meta-analysis. J Pediatr Psychol 2007, 32:527–541.

48. Shillingford AJ, Glanzman MM, Ittenbach RF, Clancy RR, Gaynor JW, Wernovsky G: Inattention, hyperactivity, and school performance in a population of school-age children with complex congenital heart disease. Pediatrics 2008, 121:e759–767.

49. Kovacs AH, Saidi AS, Kuhl EA, Sears SF, Silversides C, Harrison JL, Ong L, Colman J, Oechslin E, Nolan RP: Depression and anxiety in adult congenital heart disease: predictors and prevalence. Int J Cardiol 2009, 137:158–164.

50. Bishop SL, Thurm A, Farmer C, Lord C: Autism Spectrum Disorder, Intellectual Disability, and Delayed Walking. Pediatrics 2016, 137:e20152959.

51. De Rubeis S, He X, Goldberg AP, Poultney CS, Samocha K, Cicek AE, Kou Y, Liu L, Fromer M, Walker S, et al: Synaptic, transcriptional and chromatin genes disrupted in autism. Nature 2014, 515:209–215.

52. Noterdaeme M, Minow F, Amorosa H: [Applicability of the Child Behavior Checklist in developmentally delayed children]. Z Kinder Jugendpsychiatr Psychother 1999, 27:183–188.

53. Mazefsky CA, Anderson R, Conner CM, Minshew N: Child Behavior Checklist Scores for School-Aged Children with Autism: Preliminary Evidence of Patterns Suggesting the Need for Referral. Journal of psychopathology and behavioral assessment 2011, 33:31–37.

54. Bruining H, Eijkemans MJ, Kas MJ, Curran SR, Vorstman JA, Bolton PF: Behavioral signatures related to genetic disorders in autism. Mol Autism 2014, 5:11.

55. Klein-Tasman BP, Mervis CB: Autism Spectrum Symptomatology Among Children with Duplication 7q11.23 Syndrome. J Autism Dev Disord 2018, 48:1982–1994.

56. Hidding E, Swaab H, de Sonneville LM, van Engeland H, Sijmens-Morcus ME, Klaassen PW, Duijff SN, Vorstman JA: Intellectual functioning in relation to autism and ADHD symptomatology in children and adolescents with 22q11.2 deletion syndrome. J Intellect Disabil Res 2015, 59:803–815.

57. Vorstman JA, Breetvelt EJ, Thode KI, Chow EW, Bassett AS: Expression of autism spectrum and schizophrenia in patients with a 22q11.2 deletion. Schizophr Res 2013, 143:55–59.

58. Vorstman JA, Morcus ME, Duijff SN, Klaassen PW, Heineman-de Boer JA, Beemer FA, Swaab H, Kahn RS, van Engeland H: The 22q11.2 deletion in children: high rate of autistic disorders and early onset of psychotic symptoms. J Am Acad Child Adolesc Psychiatry 2006, 45:1104–1113.

59. Qiao Y, Riendeau N, Koochek M, Liu X, Harvard C, Hildebrand MJ, Holden JJ, Rajcan-Separovic E, Lewis ME: Phenomic determinants of genomic variation in autism spectrum disorders. J Med Genet 2009, 46:680–688.

60. Flor J, Bellando J, Lopez M, Shui A: Developmental functioning and medical Co-morbidity profile of children with complex and essential autism. Autism Res 2017, 10:1344–1352.

61. Finlay WML, Lyons E: Methodological issues in interviewing and using self-report questionnaires with people with mental retardation. Psychological Assessment 2001, 13:319–335.

62. Fisher MH, Mello MP, Dykens EM: Who reports it best? A comparison between parent-report, self-report, and the real life social behaviors of adults with Williams syndrome. Research in Developmental Disabilities 2014, 35:3276–3284.

63. Lincoln AJ, Searcy YM, Jones W, Lord C: Social interaction behaviors discriminate young children with autism and Williams syndrome. J Am Acad Child Adolesc Psychiatry 2007, 46:323–331.

64. Riby DM, Hanley M, Kirk H, Clark F, Little K, Fleck R, Janes E, Kelso L, O’Kane F, Cole-Fletcher R, et al: The interplay between anxiety and social functioning in Williams syndrome. J Autism Dev Disord 2014, 44:1220–1229.

65. van der Fluit F, Gaffrey MS, Klein-Tasman BP: Social Cognition in Williams Syndrome: Relations between Performance on the Social Attribution Task and Cognitive and Behavioral Characteristics. Front Psychol 2012, 3:197.

66. Morris CA, Mervis CB, Paciorkowski AP, Abdul-Rahman O, Dugan SL, Rope AF, Bader P, Hendon LG, Velleman SL, Klein-Tasman BP, Osborne LR: 7q11.23 Duplication syndrome: Physical characteristics and natural history. Am J Med Genet A 2015, 167a:2916–2935.

67. Sanders SJ, Ercan-Sencicek AG, Hus V, Luo R, Murtha MT, Moreno-De-Luca D, Chu SH, Moreau MP, Gupta AR, Thomson SA, et al: Multiple recurrent de novo CNVs, including duplications of the 7q11.23 Williams syndrome region, are strongly associated with autism. Neuron 2011, 70:863–885.

68. Niarchou M, Chawner S, Doherty JL, Maillard AM, Jacquemont S, Chung WK, Green-Snyder L, Bernier RA, Goin-Kochel RP, Hanson E, et al: Psychiatric disorders in children with 16p11.2 deletion and duplication. Transl Psychiatry 2019, 9:8.

69. Zufferey F, Sherr EH, Beckmann ND, Hanson E, Maillard AM, Hippolyte L, Macé A, Ferrari C, Kutalik Z, Andrieux J, et al: A 600 kb deletion syndrome at 16p11.2 leads to energy imbalance and neuropsychiatric disorders. Journal of Medical Genetics 2012, 49:660–668.

70. Green Snyder L, D’Angelo D, Chen Q, Bernier R, Goin-Kochel RP, Wallace AS, Gerdts J, Kanne S, Berry L, Blaskey L, et al: Autism Spectrum Disorder, Developmental and Psychiatric Features in 16p11.2 Duplication. J Autism Dev Disord 2016, 46:2734–2748.

71. Nag HE, Nordgren A, Anderlid BM, Nærland T: Reversed gender ratio of autism spectrum disorder in Smith-Magenis syndrome. Mol Autism 2018, 9:1.

72. Schreiner MJ, Karlsgodt KH, Uddin LQ, Chow C, Congdon E, Jalbrzikowski M, Bearden CE: Default mode network connectivity and reciprocal social behavior in 22q11.2 deletion syndrome. Soc Cogn Affect Neurosci 2014, 9:1261–1267.

73. Niklasson L, Rasmussen P, Oskarsdottir S, Gillberg C: Autism, ADHD, mental retardation and behavior problems in 100 individuals with 22q11 deletion syndrome. Res Dev Disabil 2009, 30:763–773.

74. Sobin C, Kiley-Brabeck K, Monk SH, Khuri J, Karayiorgou M: Sex differences in the behavior of children with the 22q11 deletion syndrome. Psychiatry Res 2009, 166:24–34.

75. Zablotsky B, Black LI, Blumberg SJ: Estimated Prevalence of Children With Diagnosed Developmental Disabilities in the United States, 2014-2016. NCHS Data Brief 2017:1–8.

