## Supplemental information for "Neuropsychiatric Phenotypes and a Distinct Constellation of ASD Features in 3q29 Deletion Syndrome: Results from the 3q29 Registry"

**Table S1: Questionnaire demographics.** Characteristics of study participants with 3q29Del and controls completing each questionnaire utilized in the present study.

| | Medical and Demographic Questionnaire | | Social Responsiveness Scale (SRS) | | Social Communication Questionnaire (SCQ) | | Autism Spectrum Screening Questionnaire (ASSQ) | | Achenbach Behavior Checklists (CBCL/ABCL) | |
|---|---|---|---|---|---|---|---|---|---|---|
| | 3q29Del | Control | 3q29Del | Control | 3q29Del | Control | 3q29Del | Control | 3q29Del | Control |
| Age, years (mean ± SD) | 10.0 ± 8.6 | 9.9 ± 7.2 | 10.7 ± 7.6 | 10.4 ± 7.3 | 11.2 ± 6.5 | 11.8 ± 6.9 | 10.4 ± 3.9 | 10.9 ± 3.7 | 9.8 ± 6.8 | 9.5 ± 7.0 |
| Sex (%, n) | | | | | | | | | | |
| Male | 58.1% (54) | 51.6% (33) | 54.2% (26) | 53.6% (30) | 57.6% (19) | 52.2% (24) | 66.7% (16) | 48.6% (17) | 58.3% (28) | 49.1% (28) |
| Female | 41.9% (39) | 48.4% (31) | 45.8% (22) | 46.4% (26) | 42.4% (14) | 47.8% (22) | 33.3% (8) | 51.4% (18) | 41.7% (20) | 50.9% (29) |

**Sensitivity analysis testing for effect of self-registrants**

To examine the effect of self-registrants on all analyses, we stratified participants based on registration status. Of 93 total 3q29Del cases, 3 were self-registered (3.2%) and 90 were parent-registered (96.8%). Of 64 total controls, 0 were self-registered (0%) and 64 were parent-registered (100%). Self-registrants in the study population were considered to have a significant effect on the results if the estimates for any analysis were changed by 10% or more, or if the conclusions from any analyses changed. All analyses were conducted identically when comparing the complete data and the stratified data, as outlined in Methods.

**Table S2: Sensitivity analysis for self-reposted neuropsychiatric diagnoses**. Comparison of self-reported diagnoses in the full 3q29Del dataset versus the reduced dataset with self-registrants removed. Asterisks indicate where 3q29Del was compared to general population prevalence values [41, 75]. P values were calculated with one-sample proportion test with Yates' continuity correction when comparing to population prevalence and Fisher's exact test when comparing to controls.

| | | 3q29Del Prevalence | P value; 3q29Del vs. Control |
|---|---|---|---|
| GDD/MR* | | | |
| | Full dataset | 55 (59.1%) | <2.2E-16 |
| | Reduced dataset | 55 (61.1%) | <2.2E-16 |
| ASD* | | | |
| | Full dataset | 27 (29.0%) | <2.2E-16 |
| | Reduced dataset | 26 (28.9%) | <2.2E-16 |
| Anxiety | | | |
| | Full dataset | 26 (28.0%) | 0.0007 |
| | Reduced dataset | 25 (27.8%) | 0.0007 |
| Bipolar/Manic Depression | | | |
| | Full dataset | 4 (4.3%) | 0.146 |
| | Reduced dataset | 3 (3.3%) | 0.267 |
| Conduct Disorder | | | |
| | Full dataset | 1 (1.1%) | 1.00 |
| | Reduced dataset | 1 (1.1%) | 1.00 |
| Depression | | | |
| | Full dataset | 6 (6.5%) | 1.00 |
| | Reduced dataset | 5 (5.6%) | 1.00 |
| Oppositional Defiant Disorder | | | |
| | Full dataset | 3 (3.2%) | 0.271 |
| | Reduced dataset | 3 (3.3%) | 0.267 |
| Panic Attacks | | | |
| | Full dataset | 12 (12.9%) | 0.045 |
| | Reduced dataset | 12 (13.3%) | 0.044 |
| Schizophrenia | | | |
| | Full dataset | 4 (4.3%) | 0.144 |
| | Reduced dataset | 4 (4.4%) | 0.142 |

**Table S3: Sensitivity analysis for symptomology questionnaire scores.** Comparison of estimates and p values for the contribution of the 3q29 deletion versus controls to scores on the SRS, SCQ, ASSQ, and CBCL/ABCL for the full dataset and the reduced dataset with self-registrants removed. P values were calculated using simple linear regression, adjusting for age, race, and sex.

| | | Estimate | P value |
|---|---|---|---|
| SRS | | | |
| | Full dataset | 26.38 | <2.00E-16 |
| | Reduced dataset | 26.35 | <2.00E-16 |
| SCQ | | | |
| | Full dataset | 10.71 | 2.40E-13 |
| | Reduced dataset | 10.80 | 3.44E-13 |
| ASSQ | | | |
| | Full dataset | 19.61 | 2.19E-12 |
| | Reduced dataset | 19.61 | 2.19E-12 |
| CBCL/ABCL | | | |
| | Full dataset | 20.58 | 4.30E-16 |
| | Reduced dataset | 20.62 | 7.55E-16 |

Self-registrants did not have a significant effect on the conclusions from this study; none of the estimates changed by more than 10%, and the conclusion from each analysis was consistent for the full and reduced datasets.

**Table S4: Symptomology questionnaire score comparison.** Comparison of mean scores on each symptom questionnaire between study participants with 3q29Del and controls. P values were calculated using simple linear regression, adjusting for age, race, and sex.

| Scale | 3q29Del (mean ± SD) | Control (mean ± SD) | P value |
|---|---|---|---|
| SRS | 71.8 ± 14.6 | 45.9 ± 8.0 | <2.00E-16 |
| SCQ | 13.9 ± 7.4 | 3.5 ± 3.1 | 2.40E-13 |
| ASSQ | 22.2 ± 11.4 | 2.2 ± 3.4 | 2.19E-12 |
| CBCL/ABCL | 62.5 ± 10.4 | 41.8 ± 9.8 | 4.30E-16 |

**Table S5: SRS score comparison stratified by ASD status and sex.** Comparison of mean scores on the SRS between study participants with 3q29Del stratified by ASD status and sex to controls (mean ± SD = 45.9 ± 8.0). P values were calculated using simple linear regression, adjusting for age.

| | Mean ± SD | P value |
|---|---|---|
| Sex | | |
| Male control | 45.80 ± 6.38 | - |
| Male 3q29Del | 74.31 ± 15.97 | 7.70E-11 |
| Female control | 46.04 ± 9.62 | - |
| Female 3q29Del | 68.73 ± 12.40 | 7.42E-09 |
| ASD Status | | |
| Control | 45.91 ± 7.97 | - |
| No ASD diagnosis 3q29Del | 65.90 ± 13.87 | 2.16E-13 |
| ASD diagnosis 3q29Del | 82.41 ± 8.70 | <2.00E-16 |
| ASD Status and Sex | | |
| Male control | 45.80 ± 6.38 | - |
| Male 3q29Del, no ASD diagnosis | 66.29 ± 16.17 | 4.33E-07 |
| Male 3q29Del, ASD diagnosis | 83.67 ± 9.63 | 1.13E-13 |
| Female control | 46.04 ± 9.62 | - |
| Female 3q29Del, no ASD diagnosis | 65.59 ± 12.17 | 1.60E-07 |
| Female 3q29Del, ASD diagnosis | 79.40 ± 5.59 | 3.07E-08 |

**Table S6: SCQ score comparison stratified by ASD status and sex.** Comparison of mean scores on the SCQ between study participants with 3q29Del stratified by ASD status and sex to controls (mean ± SD = 3.5 ± 3.1). P values were calculated using simple linear regression, adjusting for age.

| | | Mean ± SD | P value |
|---|---|---|---|
| Sex | | | |
| | Male control | 3.38 ± 2.14 | - |
| | Male 3q29Del | 16.00 ± 8.33 | 7.64E-08 |
| | Female control | 3.68 ± 3.97 | - |
| | Female 3q29Del | 11.00 ± 4.76 | 4.44E-07 |
| ASD Status | | | |
| | Control | 3.52 ± 3.12 | - |
| | No ASD diagnosis 3q29Del | 11.16 ± 6.53 | 2.17E-08 |
| | ASD diagnosis 3q29Del | 17.57 ± 7.05 | 3.78E-14 |
| ASD Status and Sex | | | |
| | Male control | 3.38 ± 2.14 | - |
| | Male 3q29Del, no ASD diagnosis | 12.33 ± 8.25 | 1.67E-04 |
| | Male 3q29Del, ASD diagnosis | 19.30 ± 7.27 | 1.19E-08 |
| | Female control | 3.68 ± 3.97 | - |
| | Female 3q29Del, no ASD diagnosis | 10.10 ± 4.72 | 1.65E-05 |
| | Female 3q29Del, ASD diagnosis | 13.25 ± 4.65 | 6.71E-06 |

**Table S7: CBCL/ABCL DSM-oriented sub-scale score comparison.** Comparison of mean scores on the CBCL/ABCL DSM-oriented attention deficit/hyperactivity problems, anxiety problems, and depressive problems sub-scales between 3q29Del participants and controls. P values were calculated using simple linear regression, adjusting for age, race, and sex.

| DSM-oriented sub-scale | 3q29Del (mean ± SD) | Control (mean ± SD) | P value |
|---|---|---|---|
| Attention deficit/hyperactivity Problems | 60.96 ± 8.17 | 51.30 ± 2.63 | 4.70E-13 |
| Anxiety Problems | 60.94 ± 10.67 | 52.93 ± 5.95 | 1.74E-05 |
| Depressive Problems | 62.65 ± 8.41 | 52.28 ± 5.69 | 1.65E-11 |

**Table S8: Contribution of congenital heart defects to phenotypes of interest.** Examination of the relationship between congenital heart defects and self-reported ASD and GDD/MR diagnoses and age at walking within 3q29Del cases. P values were calculated using logistic (ASD, GDD/MR) and ordinal (age at walking) regressions.

| Outcome | Covariate | Estimate | P value |
|---|---|---|---|
| ASD | | | |
| | Heart defect | 0.02 | 0.967 |
| | Sex | 0.79 | 0.152 |
| | Age | 0.03 | 0.318 |
| | Race | 1.25 | 0.259 |
| GDD/MR | | | |
| | Heart defect | -0.52 | 0.278 |
| | Sex | -0.005 | 0.992 |
| | Age | -0.0006 | 0.984 |
| | Race | 0.46 | 0.513 |
| Age at walking | | | |
| | Heart defect | 0.25 | 0.648 |
| | Sex | 0.22 | 0.641 |
| | Age | -0.04 | 0.291 |
| | Race | -0.24 | 0.733 |

**Table S9: Test for confounding factors contributing to symptomology questionnaire scores.** Possible confounding factors for the increased symptom questionnaire scores observed in 3q29Del participants. With the exception of the presence of heart defects being significantly associated with the SCQ and CBCL/ABCL scores, no other confounders were significantly associated with scores. P values were calculated using simple linear models; separate models were run for each predictor-scale pair, with the exception of the age at walking variable (comparing both "delayed" and "extremely delayed" to "normal" within the same model). All models were run controlling for age, race, and sex

| Scale | Predictor | Estimate | P value |
|---|---|---|---|
| SRS | | | |
| | Heart defect | 6.66 | 0.200 |
| | GDD/MR | 6.72 | 0.135 |
| | Age at walking (Delayed) | 3.66 | 0.504 |
| | Age at walking (Extremely delayed) | 6.83 | 0.373 |
| SCQ | | | |
| | Heart defect | 6.93 | 0.039 |
| | GDD/MR | 0.61 | 0.835 |
| | Age at walking (Delayed) | -0.56 | 0.867 |
| | Age at walking (Extremely delayed) | 5.44 | 0.230 |
| ASSQ | | | |
| | Heart defect | 7.46 | 0.253 |
| | GDD/MR | 4.31 | 0.487 |
| | Age at walking (Delayed) | -3.52 | 0.602 |
| | Age at walking (Extremely delayed) | -2.28 | 0.799 |
| CBCL/ABCL | | | |
| | Heart defect | 7.15 | 0.043 |
| | GDD/MR | 2.91 | 0.376 |
| | Age at walking (Delayed) | 1.85 | 0.644 |
| | Age at walking (Extremely delayed) | 2.13 | 0.682 |

**Table S10: CBCL/ABCL Withdrawn sub-scale score comparison.** Comparison of mean scores on the CBCL/ABCL Withdrawn sub-scale between 3q29Del participants and controls (mean ± SD = 52.3 ± 5.8). 3q29Del participants were stratified by ASD status and sex for further analysis. P values were calculated using simple linear regression, adjusting for age, race, and sex.

| | | Mean ± SD | P value |
|---|---|---|---|
| Genotype | | | |
| | Control | 52.30 ± 5.80 | - |
| | 3q29Del | 62.35 ± 8.84 | 3.11E-10 |
| Sex | | | |
| | Male control | 52.00 ± 3.74 | - |
| | Male 3q29Del | 63.04 ± 10.35 | 1.24E-05 |
| | Female control | 52.59 ± 7.33 | - |
| | Female 3q29Del | 61.40 ± 6.27 | 1.02E-05 |
| ASD Status | | | |
| | Control | 52.30 ± 5.80 | - |
| | No ASD diagnosis | 60.56 ± 7.98 | 4.39E-07 |
| | ASD diagnosis | 65.94 ± 9.62 | 3.77E-09 |

**Table S11: Heart defects present in study sample.** Types of heart defects reported by both 3q29Del and control study participants. 27 total 3q29Del participants reported heart defects: one participant reported atrial septal defect and mitral valve regurgitation; one participant reported hypoplastic left heart syndrome, interrupted aortic arch/ventricular septal defect, and single ventricle anomalies; two participants reported atrial septal defect and pulmonary valvar stenosis; one participant reported atrial septal defect, pulmonary atresia, and tricuspid valve regurgitation; one participant reported atrial septal defect and pulmonary atresia; one participant reported pulmonary valvar stenosis and ventricular septal defect; one participant reported atrial septal defect and ventricular septal defect; and one participant reported atrial septal defect, patent ductus arteriosus, pulmonary valvar stenosis, pulmonary valve regurgitation, and tricuspid valve regurgitation.

| Type of defect | 3q29Del | Control |
|---|---|---|
| Aortic valvar stenosis | 2 | 0 |
| Atrial septal defect | 7 | 0 |
| Atrioventricular septal defect (or atrioventricular canal defect) | 1 | 0 |
| Hypoplastic left heart syndrome | 1 | 0 |
| Interrupted aortic arch/Ventricular septal defect | 1 | 0 |
| Mitral valve regurgitation | 1 | 0 |
| Patent ductus arteriosus | 8 | 0 |
| Pulmonary atresia | 3 | 0 |
| Pulmonary valvar stenosis | 5 | 0 |
| Pulmonary valve regurgitation | 1 | 0 |
| Single ventricle anomalies | 1 | 0 |
| Tricuspid valve regurgitation | 2 | 0 |
| Ventricular septal defect | 2 | 0 |
| Unsure/Not indicated | 6 | 2 |

**Table S12: Comparison of 3q29 registry-leveraged and gold-standard phenotyping measures.** SRS total scores and category (Normal, Mild, Moderate, Severe) for 16 (56.25% male) study participants with 3q29Del, with parent-reported ASD diagnosis in the Medical & Demographic Questionnaire via the online 3q29 registry, and ASD diagnosis as determined by gold-standard direct evaluation by members of the Emory 3q29 Project team. Note that all parent-reported ASD diagnoses are supported by direct, in-person phenotyping, with one participant (3558) qualifying for a new ASD diagnosis after gold-standard phenotyping.

| Subject ID | Sex | Age (years) | SRS total score | SRS category | Parent-reported ASD diagnosis | Gold-standard ASD diagnosis |
|---|---|---|---|---|---|---|
| 3557 | Female | 17.5 | 83 | Severe | Yes | Yes |
| 3563 | Female | 21.17 | 73 | Moderate | Yes | Yes |
| 3548 | Male | 14 | 70 | Moderate | Yes | Yes |
| 3558 | Male | 10.83 | 90 | Severe | No | Yes |
| 3678 | Male | 16.08 | 81 | Severe | Yes | Yes |
| 3600 | Female | 10.5 | 56 | Normal | No | No |
| 3607 | Female | 8.67 | 63 | Mild | No | No |
| 3627 | Female | 6.08 | 78 | Severe | No | No |
| 3658 | Female | 27.33 | 54 | Normal | No | No |
| 3797 | Female | 14.92 | 50 | Normal | No | No |
| 3540 | Male | 7.67 | 69 | Moderate | No | No |
| 3575 | Male | 15.83 | 84 | Severe | No | No |
| 3582 | Male | 18.17 | 66 | Moderate | No | No |
| 3590 | Male | 7.42 | 56 | Normal | No | No |
| 3625 | Male | 18.67 | 82 | Severe | No | No |
| 3647 | Male | 9 | 42 | Normal | No | No |

**Table S13: Comparison of CBCL/ABCL sub-scale scores between 3q29Del and 22q11.2 deletion.** Comparison of the mean scores for 3q29Del on the CBCL/ABCL Withdrawn, Somatic Complaints, Anxious/Depressed, Social Problems, and Thought Problems sub-scales to those reported in a sample of 22q11.2 deletion probands [71]. The Social Problems sub-scale is only included on the school-age CBCL; the Thought Problems sub-scale is included on the school-age CBCL and ABCL. 3q29Del sample size for each sub-scale is indicated in parentheses.

| CBCL/ABCL sub-scale | 3q29Del mean (n) | 22q11.2 deletion mean | P value |
|---|---|---|---|
| Withdrawn | 62.4 (48) | 58.2 | 0.0010 |
| Somatic Complaints | 62.0 (48) | 57.4 | 0.0002 |
| Anxious/Depressed | 60.3 (48) | 57.2 | 0.0157 |
| Social Problems | 65.5 (26) | 62.4 | 0.0053 |
| Thought Problems | 67.9 (31) | 61.1 | 0.0004 |

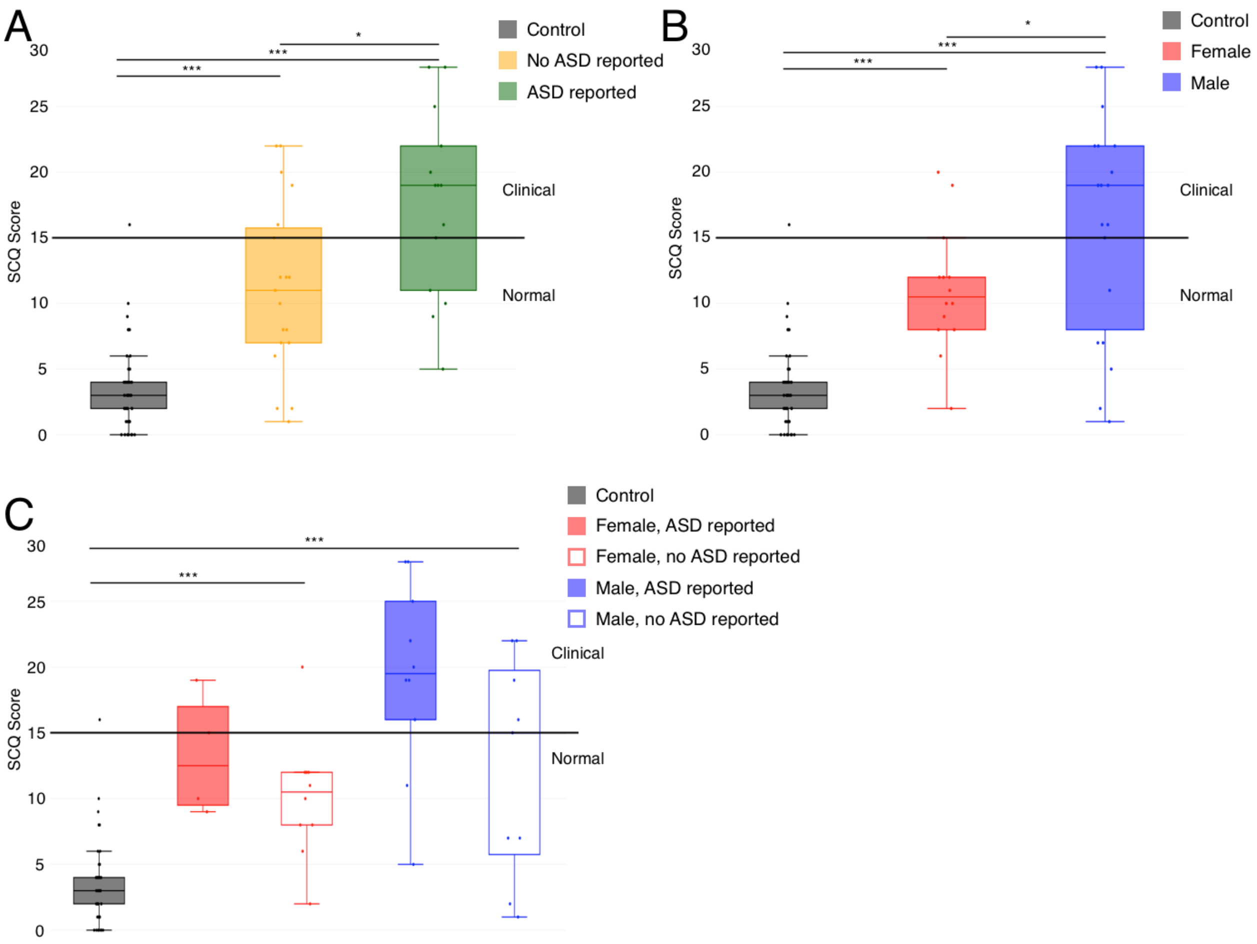


**Figure S1. A)** SCQ scores split by control (n=46), 3q29Del not reporting an ASD diagnosis (n=19), and 3q29Del reporting an ASD diagnosis (n=14), showing a significant association between self-reported diagnostic status and SRS score. **B)** SCQ scores split by sex, with control (n=46), 3q29Del female (n=14), and 3q29Del male (n=19), showing a lack of sex bias in scores for 3q29Del participants. **C)** SCQ scores split by sex and self-reported diagnostic status, with control (n=46), 3q29Del female reporting ASD (n=4), 3q29Del female not reporting ASD (n=10), 3q29Del male reporting ASD (n=10), and 3q29Del male not reporting ASD (n=9), showing inflated scores for 3q29Del participants irrespective of sex or diagnostic status. *, $p < 0.05$; ***, $p < 0.001$

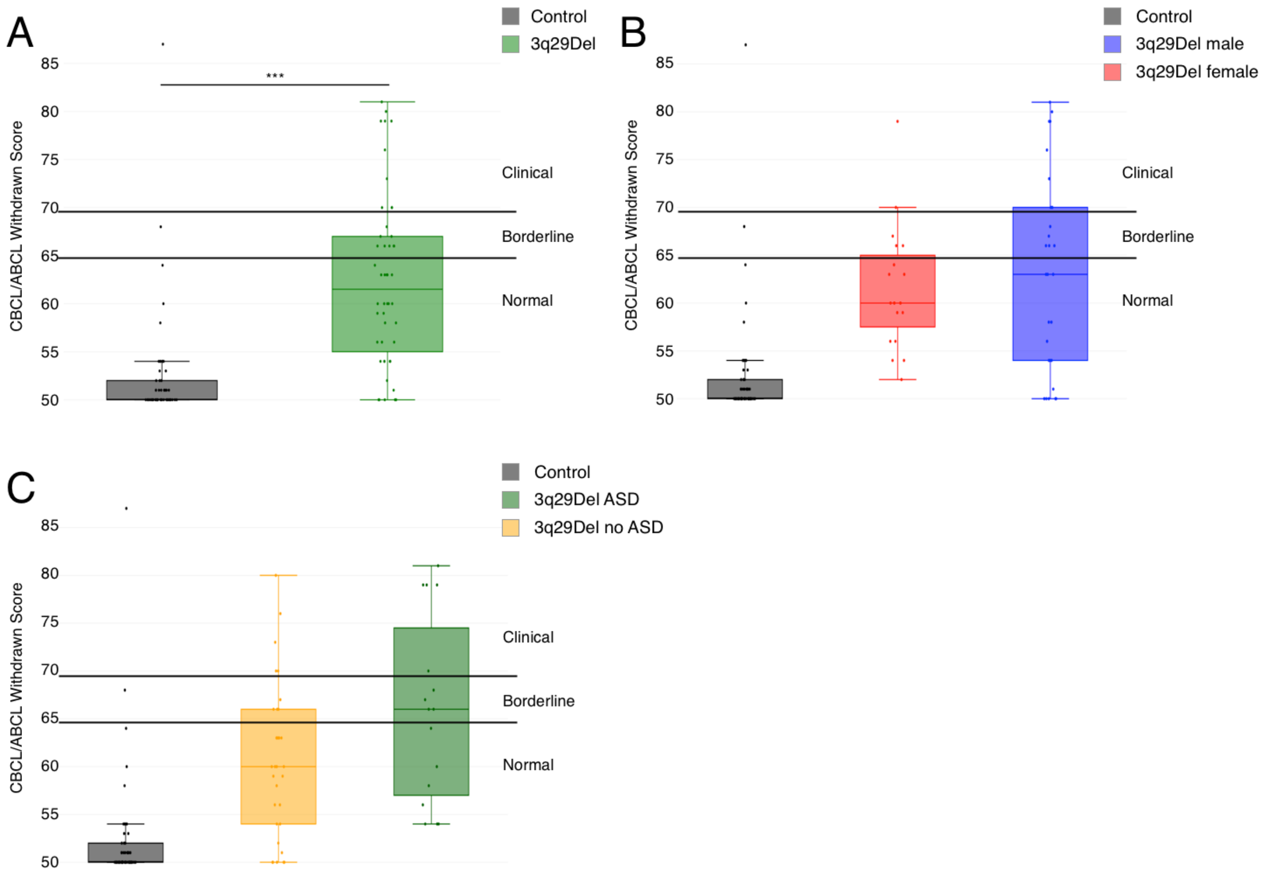


**Figure S2. A)** Profile of 3q29Del participants (n=48) and controls (n=57) on the Withdrawn sub-scale from the CBCL and ABCL, showing a significantly higher score in 3q29Del participants, with a mean score for both groups in the normal range. **B)** Profile of 3q29Del males (n=28) and females (n=20) and controls (n=57) on the Withdrawn sub-scale from the CBCL and ABCL, showing that scores are not significantly different between males and females. **C)** Profile of 3q29Del participants reporting an ASD diagnosis (n=16) and not reporting an ASD diagnosis (n=32) and controls (n=57) on the Withdrawn sub-scale from the CBCL and ABCL, showing that scores are not significantly different for 3q29Del participants based on self-reported ASD status. ***, $p < 0.001$

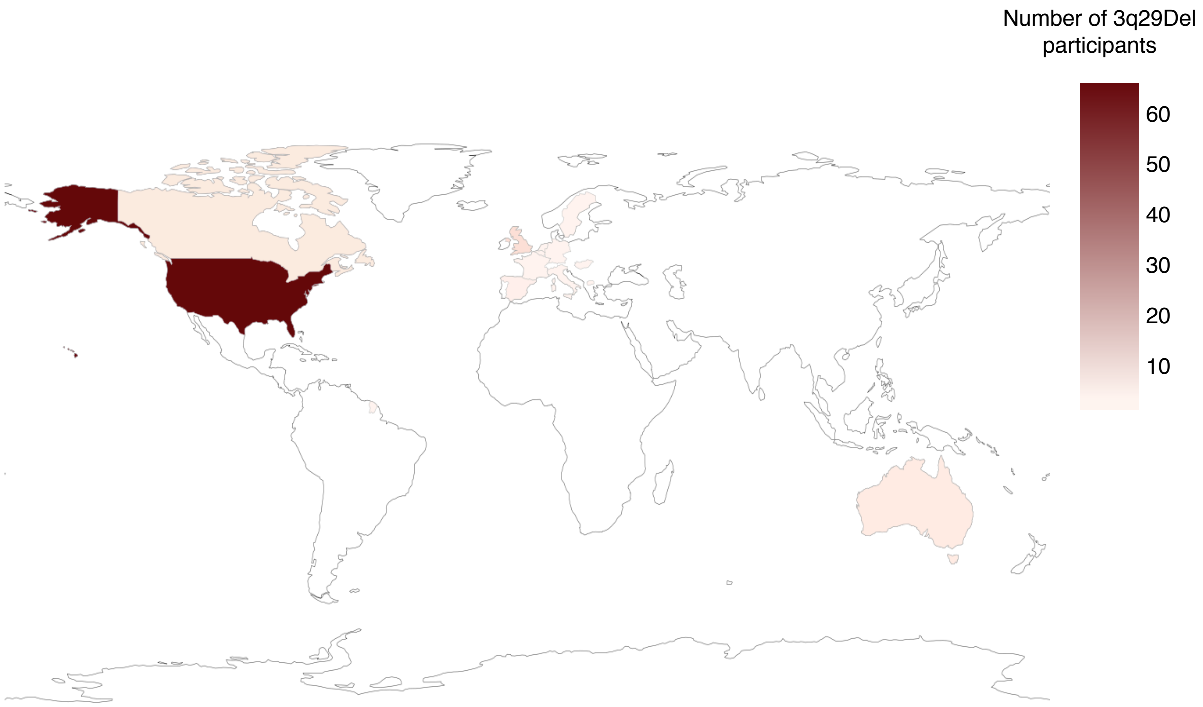


**Figure S3.** Geographic distribution of participants with 3q29Del (n=93). White indicates countries not represented in the present study sample.
